# SUBATOMIC: a SUbgraph BAsed mulTi-OMIcs Clustering framework to analyze integrated multi-edge networks

**DOI:** 10.1101/2022.06.01.494279

**Authors:** Jens Uwe Loers, Vanessa Vermeirssen

## Abstract

Representing the complex interplay between different types of biomolecules across different omics layers in multi-omics networks bears great potential to gain a deep mechanistic understanding of gene regulation and disease. However, multi-omics networks easily grow into giant hairball structures that hamper biological interpretation. Module detection methods can decompose these networks into smaller interpretable modules. However, these methods are not adapted to deal with multi-omics data nor consider topological features. When deriving very large modules or ignoring the broader network context, interpretability remains limited. To address these issues, we developed a subgraph based multi-omics clustering framework (SUBATOMIC), which infers small and interpretable modules with a specific topology while keeping track of connections to other modules and regulators.

SUBATOMIC groups specific molecular interactions in composite network subgraphs of two and three nodes and clusters them into topological modules. These are functionally annotated, visualized and overlaid with expression profiles to go from static to dynamic modules. To preserve the larger network context, SUBATOMIC investigates statistically the connections in between modules as well as between modules and regulators such as miRNAs and transcription factors. We applied SUBATOMIC to analyze a composite *Homo sapiens* network containing transcription factor-target gene, miRNA-target gene, protein-protein, homologous and co-functional interactions from different databases. We derived and annotated 5586 modules with diverse topological, functional and regulatory properties. We created novel functional hypotheses for unannotated genes. Furthermore, we integrated modules with condition specific expression data to study the influence of hypoxia in three cancer cell lines. We developed two prioritization strategies to identify the most relevant modules in specific biological contexts: one considering GO term enrichments and one calculating an activity score reflecting the degree of differential expression. Both strategies yielded modules specifically reacting to low oxygen levels.

In conclusion, we developed the SUBATOMIC framework that generates interpretable modules from multi-omics networks and applied it to hypoxia in cancer. SUBATOMIC can infer and contextualize modules, explore condition or disease specific modules, identify regulators and functionally related modules, and derive novel gene functions for uncharacterized genes. The software is available at https://github.com/CBIGR/SUBATOMIC.

## Introduction

Eukaryotic gene regulation involves a complex interplay between different types of biomolecules to safeguard correct gene expression in space and time. Transcription factors (TFs) bind to specific sequences in the DNA such as promoter and enhancer regions to activate or repress gene expression [1–3]. Co-factors bind to TFs and interact with the transcriptional machinery [2, 4]. At the epigenetic level, the accessibility of chromatin is the degree to which molecules such as TFs, RNA-polymerases or chromatin organizing proteins are able to establish a physical contact with the underlying DNA via promotor, enhancer and insulator regions [5]. The accessibility is dynamic and changes in response to external stimuli as well as developmental signals leading to notable differences in expression between various cell types [5, 6]. Several classes of non-coding RNA (ncRNA) also have an impact on gene regulation. Micro-RNAs (miRNA) suppress protein translation or induce messenger RNA (mRNA) degradation, mostly by binding to the 3’-UTR of target messenger RNAs [7, 8]. Moreover, they are regulated by DNA methylation, histone modifications and more than 140 forms of RNA modifications [9]. In turn, miRNAs themselves target epigenetic-associated enzymes such as DNA methyltransferases, ten-eleven translocation genes and histone deacetylases [9, 10]. Long non-coding RNAs (lncRNAs) act as signal molecules that mediate transcription of downstream genes, as decoy molecules to repress biological processes and pathways by binding TFs and blocking their regulatory activity [11, 12] or compete with mRNAs for miRNA binding [13, 14]. Additionally, several genes, especially regulatory proteins such as TFs and miRNAs, have undergone duplication events during their evolution, leading to gene redundancy or/and the acquisition of novel biological functions over time [15, 16].

High-throughput technologies, like RNA-seq, ChIP-seq and mass spectrometry yield an enormous amount of high-quality data in the context of gene regulation. These data are available in databases that continuously grow by adding and integrating novel datatypes and datasets. One example is the resource ‘Discriminant Regulon Expression Analysis’ (DORothEA) [17, 18]. It contains signed TF-target interactions based on literature-curated resources, ChIP-seq peaks, gene expression based inference and TF binding sites information [17, 18]. DORotheEA is embedded in the OmniPath database, which includes many additional interaction types such as miRNA-target interactions, lncRNA-target interactions, ligand- receptor binding and protein-protein interactions [19, 20]. HumanNet is a human gene network resource that captures co-functional and physically binding interactions: the co-functional network (COF) includes co-essentiality and co-expression interactions, while the protein- protein interaction network contains literature-curated and high-throughput interactions of physically binding proteins [21]. Many more databases exist for diverse types of molecular interactions and their size and number continuously grow.

To understand gene regulation in depth, we need to comprehend how different molecular interactions together coordinate phenotype-specific gene expression. Indeed, several studies have shown that considering complementary molecular interactions increases our understanding of regulatory processes. Co-expressed genes and genes encoding physically interacting proteins are often regulated by the same set of TFs [22, 23]. Genes encoding TFs that control miRNA expression have a higher chance to be post-transcriptionally repressed by the miRNA [24]. Genes co-regulated by miRNAs show weaker functional links compared to TF-regulated genes [25]. These complex, diverse interactions between several biomolecules in gene regulation can be modelled at a systems level in gene regulatory networks (GRNs). GRNs map the molecular interactions between regulators, mainly TFs, and their target genes, based on relevant high-throughput data, with or without using computational inference [26, 27]. Integrated GRNs take into account different types of molecular interactions implicated in gene regulation [28]. Currently, proficient methods for integrating multi-omics data into these GRNs are still lacking, as well as methods for the analysis and biological interpretation of intricate, integrated networks.

Biological networks are often hard to interpret as a whole. They possess a high number of nodes and edges merged into a giant ‘hairball’ structure that makes a meaningful visualization and their functional interpretation extremely challenging [29, 30]. Many approaches have been developed to tackle this problem. Their shared principle is to decompose those hairball structure into smaller interpretable subnetworks, often referred to as modules or communities. Methods can be co-expression based, topology based, pan-sample based and multi-edge based including tools such as WGCNA, SimMod, ModulOmics, and LemonTree [31–41]. WGCNA clusters genes with high expression correlation and summarizes modules using their module eigengene (Azad, 2017; Azad and Lee, 2013). SimMod uses a mixed integer non-linear programming model to integrate WGCNA based co-expression networks with physical and genetic interactions into multi-omics communities [33]. ModulOmics identifies de-novo cancer driver pathways and modules by integrating protein-protein interactions, mutual exclusivity of mutations and copy number variations (CNV), transcriptional co-regulation and co-expression [40]. Other methods integrated TF-target gene interactions and protein-protein interactions with a ‘function-to-structure’ based method by deriving modules based on genes with a shared GO-annotation [41]. LemonTree tries to bridge functions from different approaches by first inferring a set of co-expression based modules across different datasets, second integrating them in a consensus module and finally integrates these modules with potential multi-omics regulators for example TF, miRNAs or CNV [36]. While existing approaches strongly increased the interpretability of large multi-omics networks, some challenges remain. One common limitation is that the number of derived clusters is usually very small and contains a lot of genes. Most large modules correlate well with biological properties or phenotypes but lack detailed and causal interpretation. Moreover, modules are interpreted as completely separated entities and do not share any genes. However, when considering topology in multi-edge networks, genes can appear in different topological contexts and thus in different modules. On the other hand, given that many small and interpretable modules exist, keeping track of the inter-module relationships is crucial to not miss out on a broader network interpretation. Thus a method that considers network topology, edge-causality and condition specific data (e.g. expression) while producing small and interpretable modules in a specific network context can substantially complement the existing inference methods.

We previously proposed a data integration framework for the worm *Caenorhabditis elegans* and the plant *Arabidopsis thaliana* that groups specific molecular interactions in composite network subgraphs, clusters these next into biologically relevant, topological modules, highlights connections between modules and regulators, and finally overlays these modules with gene expression profiles to go from static to dynamic modules [28]. We learned that different molecular interactions interrelate in distinct topologic modules with specific biological functions to generate a coordinated response in gene regulation. Here, we extended this data integration framework to SUbgraph BAsed mulTi-OMIcs Clustering (SUBATOMIC). SUBATOMIC infers composite subgraph-based modules from diverse interaction databases and gene expression profiles and analyzes them in an updated, generalized, and automated analysis framework. Upon dissecting the composite network into small interpretable modules, we keep track of interactions that connect regulators with modules as well as modules with each other in a superview to preserve their larger network context and facilitate biological interpretation. To make the static network modules dynamic and evaluate their role in specific conditions, we implemented a module activity score. The score can be used to rank modules with regard to their degree of dysregulation upon condition change. The method is applicable to any user-defined set of networks with overlapping vertices for any species of interest. We applied SUBATOMIC to integrate six networks from *H. sapiens* based on TF-target interactions, miRNA-mRNA interactions, protein-protein interactions, functional interactions, and homologous connections for proteins and miRNAs. The inferred modules allowed us to propose functional hypotheses for insufficiently annotated proteins. As proof of concept, we further contextualized our modules with expression data for cancer cell lines under hypoxic conditions. Hypoxia occurs when a cell or tissue is not sufficiently supplied with oxygen to maintain their homeostatic state (Gaspar and Velloso, 2018; Hiraga, 2018). This state frequently appears in the tumor microenvironment leading to cellular responses that increase the risk of metastasis and reduce the success of treatment [42, 43]. We identified modules sensitive to hypoxia conditions using both activity and GO term based features. We showed that these responsive modules are highly connected in our superview analysis compared with a random control. We highlighted several examples and guidelines on how to use SUBATOMIC to gain biological insights. The SUBATOMIC pipeline is available on GitHub (https://github.com/CBIGR/SUBATOMIC).

## Results

### SUBATOMIC: a subgraph based multi-omics clustering framework

We developed SUBATOMIC, a SUbgraph BAsed mulTi-Omics Clustering framework to construct and analyze multi-edge networks (Figure 1). SUBATOMIC takes networks composed of different interaction types as input. Interactions can be directed such as TF-target interactions and miRNA-target interactions or undirected such as protein-protein interactions. The networks need to have a partial overlapping node set to allow for integration over the different interaction types. Given the multi-edge networks, SUBATOMIC first uses the subgraph enumeration algorithm ISMAGS to decompose it into a set of 3-node composite subgraphs [44]. Additionally, we incorporated an own script to find specific 2-node subgraphs. Subgraphs are classified according to type and direction of edges they contain. By integrating directed and undirected edges in 2- and 3-node composite subgraphs, we identify 8 different subgraph types [28]. In a co-pointing subgraph (COP), an undirected edge connects two regulators and together they regulate a target. The co-regulated subgraph (COR) contains one regulator controlling two interacting target genes. The feed-forward loop (FFL) has a regulator that directly regulates a target gene and another regulator, which also controls the target gene. In the circular feedback subgraph (CIR), regulators act upon each other through feedback loops. The feedback undirected subgraph (FBU) consists of two directed interactions in a cascade that are connected by an undirected interaction. The feedback 2 undirected subgraph (FB2U) combines two undirected interactions and one directed interaction. The complex subgraph (COM) contains only undirected edges. Finally, the two-node feedback subgraph (2FB) couples a directed edge with an undirected edge.

**Figure 1:**
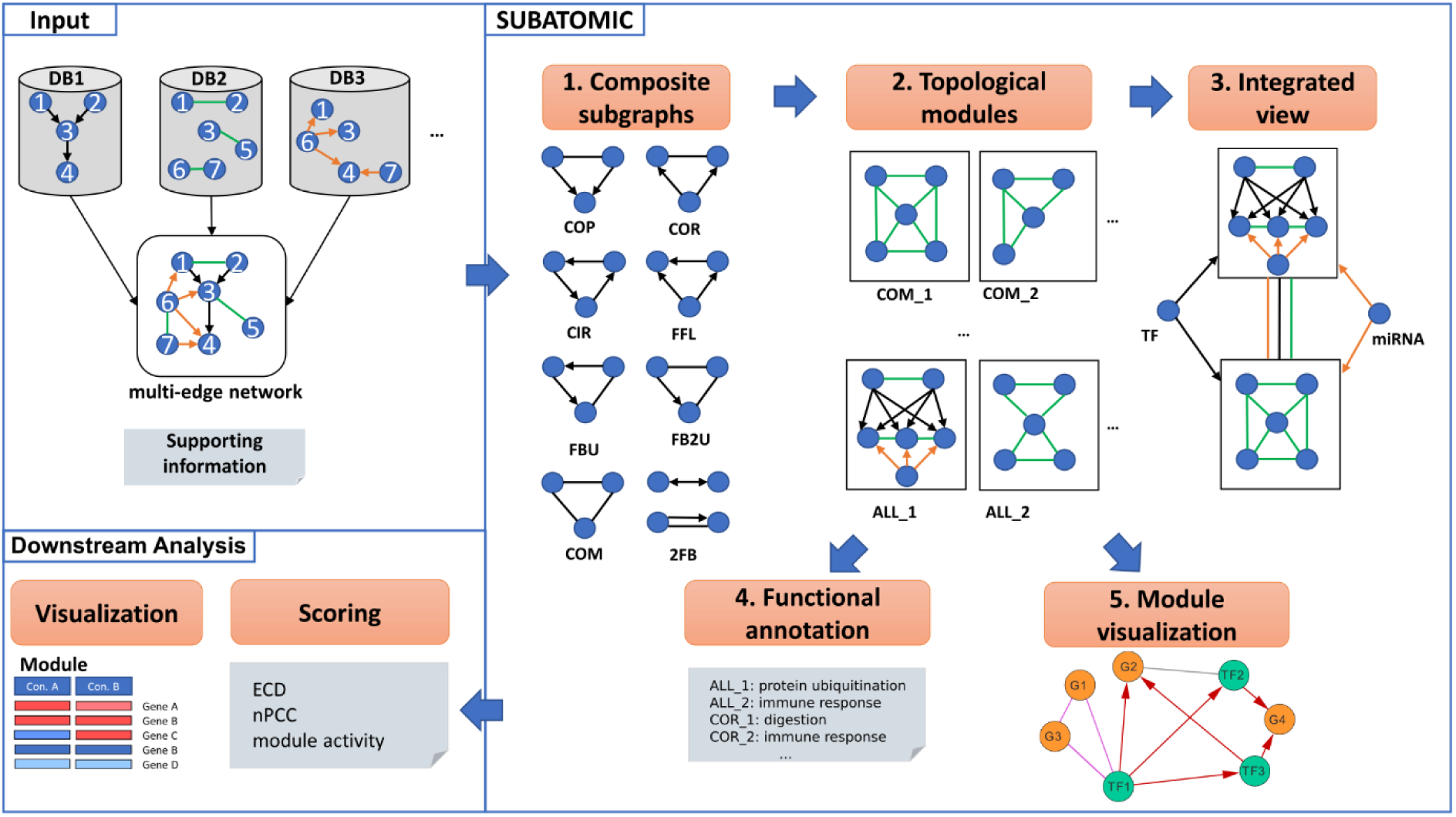
Overview of the SUBATOMIC workflow. SUBATOMIC takes as input a multi-edge network consisting of directed and/or undirected interactions of different interaction types. Supporting information contains additional files such as gene annotations or a list of subgraphs that should be considered (see methods). The multi-edge network is then decomposed into composite subgraphs using ISMAGS for co-pointing (COP), co-regulated (COR), circular (CIR), feed-forward loop (FFL), feedback undirected loop, feedback 2 undirected loop, complex (COM) and 2 feedback subgraphs. Based on the subgraphs, SCHYpe generates modules for each subgraph type as well as for all subgraph types together (ALL). Modules are connected with each other as well as with regulators to produce an integrated view (superview). Modules are functionally annotated with GOATOOLS and Cytoscape outputs for each module are generated. Networks analyzed in this way are considered static if they did not incorporate any condition specific information. In the downstream analysis, modules are integrated with condition-specific expression data. Several scores reflect the condition-specific activity of modules: the expression dynamicity score (ECD), the average Pearson correlation of expression values in a module (nPCC) and the module activity score.

We followed the ISMAGS nomenclature in representing subgraphs by their specific interaction types and edge signs [44]. Each input network of a specific interaction type is assigned a specific letter: R for TF-gene, M for miRNA-mRNA, P for protein-protein, C for co-functional and H for homologous interactions. Then each 3-node subgraph obtains a three letter representation according to the specific interaction type of its edges. For example, a PPP subgraph contains three undirected edges from the protein-protein interaction network and is hence of the COM type. RRP contains two edges from a regulatory TF-gene network and one from the protein-protein interaction network and is hence of the COR type. Subsequently, all subgraphs are clustered by the hypergraph-based spectral clustering algorithm ‘Spectral Clustering in Hypergraphs’ (SCHype) [45]. SCHype optimizes the edge-to-node ratio on hyperedges that represent the 3-node and 2-node subgraphs during clustering. The resulting modules share common topological features and possess specific biological functions [28, 45]. SUBATOMIC uses SCHype to first generate clusters within each type of subgraph (COM, CIR, FFL, …). Additionally, all subgraphs together are clustered in a module type called ‘ALL’. We further filter for subgraphs that contain between 5 and 50 genes similar to our previous approach [28]. Next, SUBATOMIC applies GOATOOLS to functionally annotate the modules based on Gene Ontology [46]. At this point, small and biologically interpretable modules have been obtained, but their network context is not yet considered. To address this, we arrange all modules in superview that connects modules with each other, and finds regulators connected to each modules. SUBATOMIC also calculates an output that can be imported in Cytoscape for module network visualization [47]. As a postprocessing step, the topological modules are integrated with expression data to study dynamics of gene regulation over different experimental conditions. In this step several metrics can be calculated to further characterize and prioritize modules in specific conditions. More details on the pipeline can be found in the methods section.

### Integrated human regulatory networks

Multi-omics data integration aids in the understanding of dysregulation in complex diseases. Our data integration framework SUBATOMIC not only makes use of multi-omics networks but also connects topological and functional information to leverage their interpretation. In this study we aimed to construct and analyze multi-omics networks for *H. sapiens* with SUBATOMIC. Therefore, we integrated TF-target gene, miRNA-mRNA, homologous, protein-protein, and co-functional interactions from public resources and finally added expression data from cancer cell lines under hypoxic conditions. We provided a layout of how SUBATOMIC can be used to investigate perturbed gene regulation in a disease context.

We included five different networks in our analysis that cover distinct interaction types that all influence gene regulation (see Table 1, Methods and Additional File 1). Two networks are directed and model regulatory relationships: the TF-target gene network (R) and the miRNA-mRNA network (M). Three networks are undirected: the homolog network (H), the protein-protein interaction network (P) and the co-functional network (C). The R network includes 53232 TF-target gene interactions from OmniPath from three different OmniPath sub-databases: DoRothEA (levels A-C), TF_target (curation score >1) and TF-miRNA interactions [19, 20]. The M network includes 11085 miRNA-mRNA interactions from OmniPath. To include gene homology, we included 10847 paralogous interactions between genes from the Ensemble archive and homologous miRNAs with identical seed sequence from miRbase [48, 49]. To include a layer of functional information we choose two mutually exclusive networks from HumanNet v2 [21, 50]. The 6637 co-functional edges contains co-essentiality, co-expression, and protein domain profile association edges. The 24773 physical protein-protein interaction network contains edges from high throughput assays such as yeast-two-hybrid and affinity purifications and from literature-curated protein-protein interactions.

**Table 1:** Overview of the different interaction types included in the multi-edge input network

| Interaction type | Letter | Directed | #Nodes | #Edges | #Regulators | #Targets | Database |
| --- | --- | --- | --- | --- | --- | --- | --- |
| Regulatory TF-gene interactions | R | Yes | 15014 | 53232 | 526 | 14488 | OmniPath |
| Regulatory miRNA-mRNA interactions | M | Yes | 4060 | 11085 | 850 | 3210 | OmniPath |
| Homologous genes | H | No | 4862 | 10847 | - | - | Ensembl |
| Protein-protein interactions | P | No | 10950 | 24773 | - | - | HumanNet |
| Co-functional interactions | C | No | 10682 | 66373 | - | - | HumanNet |

Overview of different interaction types included in the input multi-edge network for *H. sapiens*. The 'letter' column indicates the chosen letter to represent the interaction networks. Input networks were derived from three different databases and contained either directed or undirected edges. For directed edges, the number of regulators and target genes are shown.

### More than half of the detected subgraphs are interaction type specific

ISMAGS detected a total of 787347 3-node subgraphs (Figure 2). Complex subgraphs were most abundant covering 82.56% of all composite subgraphs, containing mostly CCC (413938 – 48.77%) and RRR (85435 – 10.07%), followed by PCC (80035 – 9.43%). While the overall fraction of non-COM type subgraphs might be small, rare subtypes can reveal interesting mechanistic insights. Subgraphs shared between different interaction types are less often observed than subgraphs detected within one type of interaction. A total of 534665 subgraphs contain at least two co-functional interactions, while 150582 subgraphs contain at least one co-functional and one protein-protein interaction. Subgraphs with edges from the co-functional network and the regulatory TF gene interaction network are counted with 26915 occurrences. The smallest amount of subgraphs with two edges of the same type is 7252 and comes from the miRNA-mRNA interactions, which also possess the lowest amount of shared subgraphs with the homologous network (826). While the interaction types and quantities used in Defoort et al. for *A. thaliana* and *C. elegans* were slightly different, we obtained comparable results with complex subgraphs being most abundant and subgraphs containing only protein-protein and homologous interactions having the highest subgraph counts [28].

**Figure 2.**
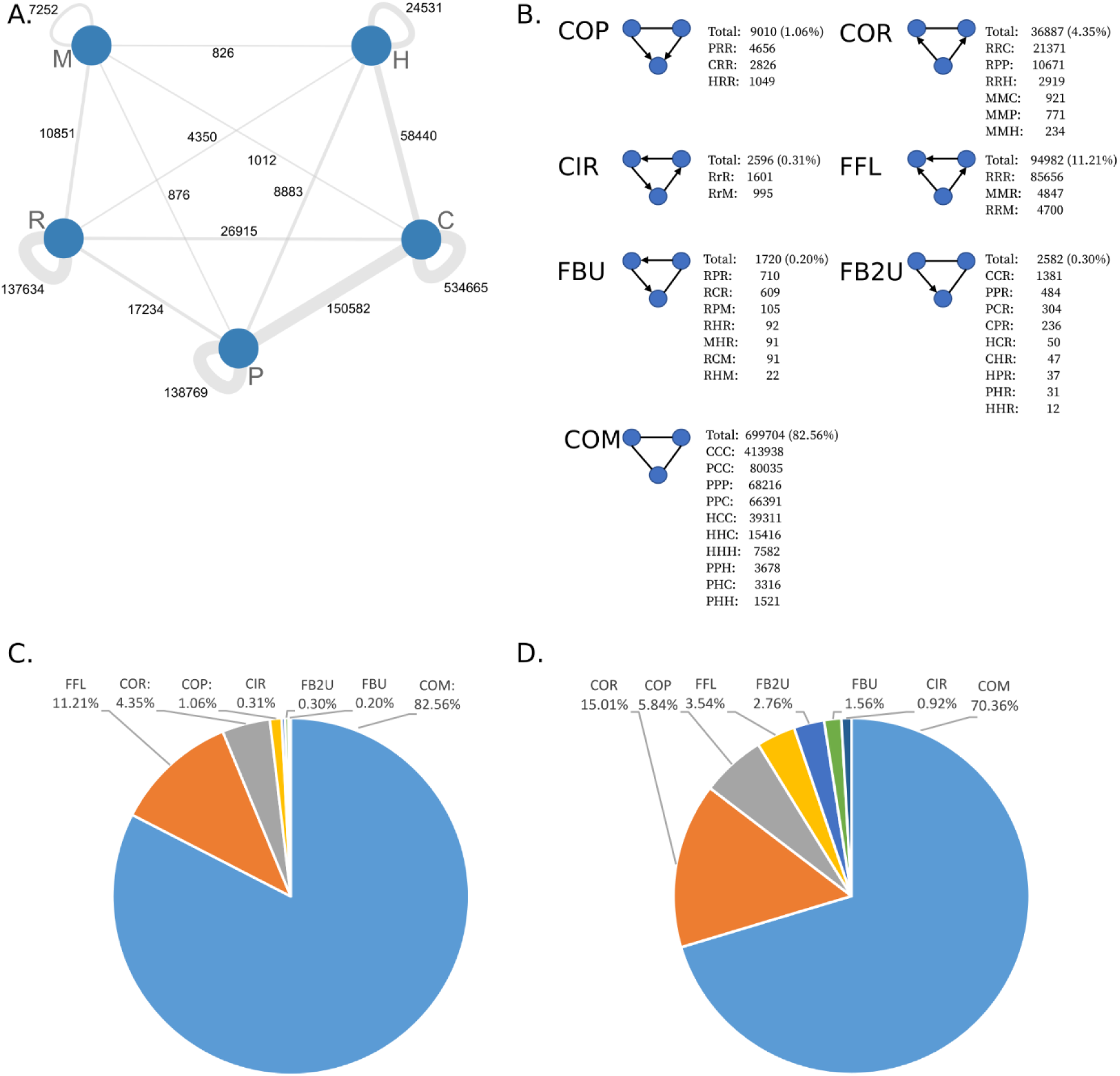
A: Interconnection between different input networks (nodes) at the number of composite subgraphs. Each edge connecting two nodes represents how many subgraphs contain at least one edge from each network. Most subgraphs contain at least two edges from the same input networks B: Overview of the counts and fractions of all different detected subgraphs. C: Overview of different subgraph types detected in the human multi-edge network. D. Overview of different module types detected in the human multi-edge network. We omitted the ALL networks in C. and D. for visualization purpose since the ALL modules contain overlapping subgraphs with other module types.

Next, SUBATOMIC uses SCHype to cluster the composite subgraphs into 7 module types. For our *H. sapiens* composite network, this yielded in a total number of 5586 modules (2762 ALL, 1987 COM, 424 COR, 165 COP, 100 FFL, 78 FB2U, 44 FBU, 26 CIR). While COM, COR, COP, FFL, FB2U, FBU and CIR are generated on mutually exclusive subgraph types, ALL contains a joint clustering of all types together and allows to find interactions between different topological modules. Hence, ALL modules were most abundant, followed by COM and COR modules. CIR modules are the least present. With regard to the context of gene regulation, COR, COP, CIR and FFL modules are the most interesting containing directed regulatory interactions.

### Most modules are densely connected and regulated by several transcription factors and miRNAs

While calculating the modules, we kept track of their larger network context in the so-called ‘superview’ analysis. This includes the interactions of modules with each other as well as with regulators such as miRNAs and TFs. We first analyzed the specificity of regulators by looking at how many modules they target. This gives insights whether regulators can be considered master regulators or specific regulators. In our analysis, we included a total of 526 TFs and 850 miRNAs.

On average, a TF targeted 6% and a miRNA targeted 2% of all modules. A total of 25 TFs regulated 5 or less modules, while five TFs are specific for only one module. Among the miRNA regulators, 90 regulated 5 or less modules while 19 only target one module. An average module is targeted by 33 TFs and 18 miRNAs. We then draw the degree distribution for in-degree and out-degree of the regulator-module interactions (Figure 3 A and B). Most modules and regulators possessed a low degree. The five highest degree modules are of FFL, CIR and ALL type. For the regulators, a high degree can be interpreted as indication for master regulators. Furthermore, we draw the distribution of the clustering coefficient for all modules (Figure 3 C). We detected a mode of 0.55-0.6 in the clustering coefficient distribution with the majority of modules showing a higher connectivity than the mode. Since our modules are strongly dominated by COM modules, excluding COM and ALL modules, now the majority of modules had a lower connectivity than the mode (Figure 3 D). As the co-functional and protein-protein interactions networks are highly interconnected, their clustering coefficient is also very high.

**Figure 3:**
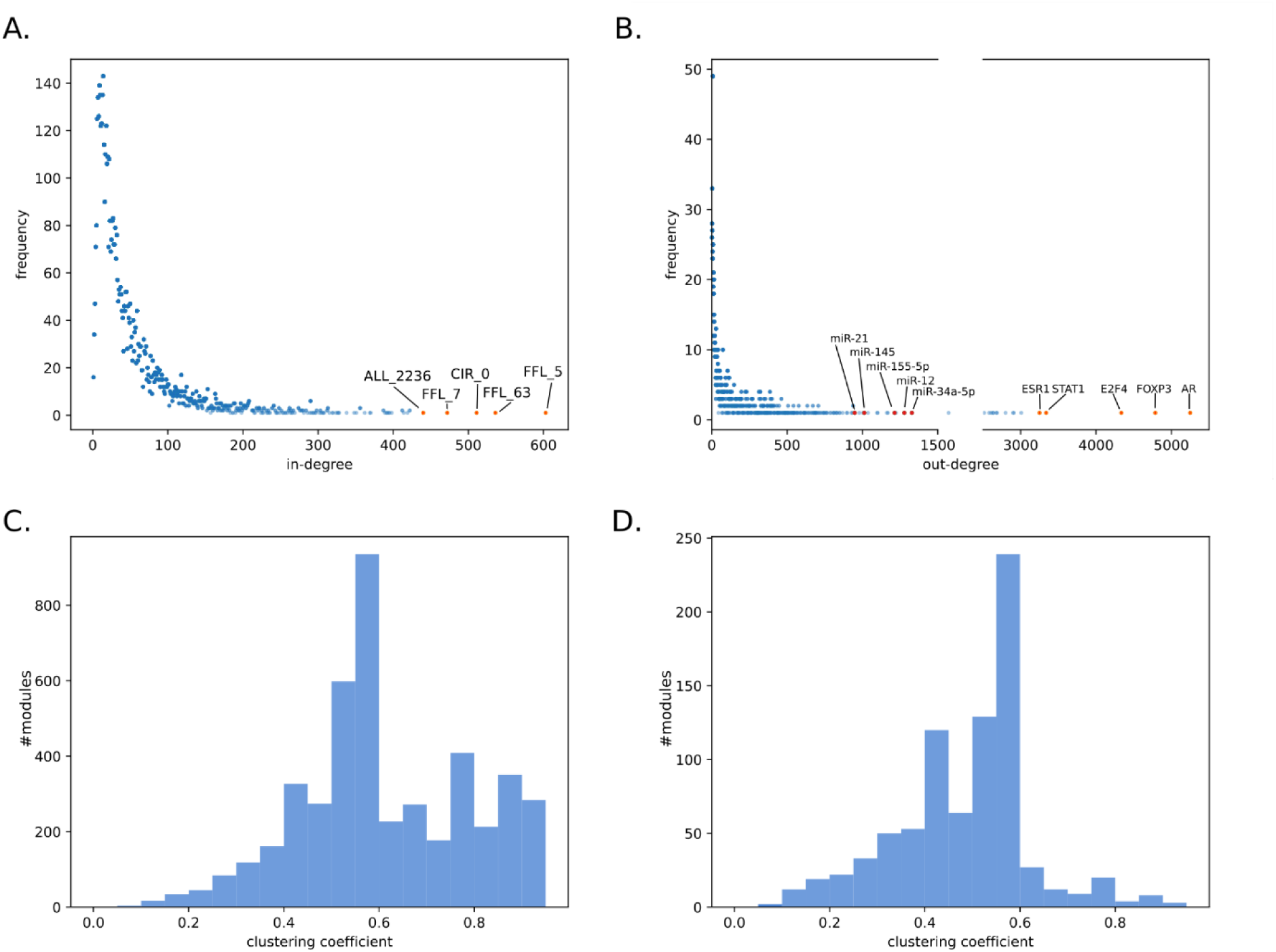
Specific network properties of the regulator-modules network. A. In-degree and B. Out-degree distribution of the interactions between regulators and modules. The top 5 modules, transcription factors and miRNAs with the highest degree are shown with label. In B., we disrupted the axis between 1500 and 2500. C. The clustering coefficient distribution of all modules. The mode was located between 0.55 and 0.6. The majority of modules had a higher connectivity than the mode. D. The clustering coefficient distribution of all modules excluding COM and ALL modules. The majority of modules had a lower connectivity than the mode. The visualization of the clustering coefficient for each independent module is given in the Additional file 6.

### GO enrichment analysis of modules reveals unknown gene functions

Upon functional GO ontology enrichment analysis, 3805 modules have a list of enriched GO terms. We can capitalize on these functional annotations by generating hypothesizes on gene functions for genes that are not well-characterized based on the guilt-by-association or guild- by-rewiring principle [51, 52]. Although most genes in our *H. sapiens* network are well-annotated and connected to many GO terms, 1404 genes have less than five GO terms and 345 have no GO term at all (Figure 4). Limiting ourselves to protein-coding genes, we found 53 genes in our modules that are annotated with merely two or less GO terms, further referenced as ‘weakly characterized genes’, 25 of which have no GO term at all. (See supplement for a complete list of un-annotated genes and their module context). Several of these genes were present in well-annotated modules and we could predict their biological function in relation to the GO annotation and structure of the module. We selected five genes without current GO annotation for further analysis: the proline and serine-rich protein 1 (PROSER1), the cyclic nucleotide-binding domain-containing protein 1 (CNBD1), the leucine, glutamate and lysine rich 1 gene (LEKR1), the RIIa domain-containing protein 1 (RIIAD1), and the glutamate-rich protein 6B (ERICH6B) (Figure 5). The gene PROSER1 (ENSG00000120685) appears in the modules ALL_1135, ALL_1880 and ALL_2888. The latter consists of a protein-protein complex of eight genes. The top 5 enriched terms are MLL3/4 complex, histone methyltransferase activity (H3-K4 specific), Set1C/COMPASS complex, histone H3-K4 methylation and histone methyltransferase complex. The histone methyltransferase complex GO term is shared by 6 out of 8 module genes. PROSER1 is directly connected to the histone methyltransferase KMT2 as well as to the PAXIP1 known to be involved in histone H3-K4 methylation [53]. Thus, we hypothesized that PROSER1 is part of a histone methylation complex. After our analysis, a recently published work confirmed the PROSER1s involvement in the regulation of various chromatin associated proteins [54]. Next, we analyzed CNBD1 (ENSG00000176571). This gene appeared in the modules ALL_654 and COM_667. The top 5 enriched terms in COM_667 are HCN channel complex, intracellular cAMP-activated cation channel activity, intracellular cyclic nucleotide activated cation channel complex, intracellular cyclic nucleotide activated cation channel activity and cyclic nucleotide- gated ion channel activity. It is connected via co-functional edges to CNGB1, CNGA1 CNGA3, and CNGA4. All four are subunits of the cyclic nucleotide gated channel (CNGA) and appear in all of the top 5 enriched terms expect for the HCN channel complex. Thus, we hypothesized that CNBD1 is also part of a cation channel complex and possesses cation channel activity. The LEKR1 (ENSG00000197980) gene appears in the modules ALL_3385 and FB2U_87. While ALL_3385 had no significant enrichment, the top 5 enriched terms in FBU_87 are cellular components muscle myosin complex, dynactin complex, myosin filament, myosin II complex and sarcomere. It is connected with MYH2, which is also annotated with all significant terms except the dynactin complex. While we cannot derive a specific function for LEKR1, we can hypothesize that it plays a role in the myosin complex. The ERICH6B (ENSG00000165837) gene appears in ALL_1153 and COM_1252. The top 5 enriched terms are the molecular functions metallocarboxypeptidase activity, carboxypeptidase activity, metalloexopeptidase activity, exopeptidase activity as well as the biological process protein processing. It has protein-protein interactions with four proteins, of which the carboxypeptidase D (CPD) is part of all enriched GO terms in this module and the succinate--CoA ligase SUCLG2 is part of the cellular amide metabolic process. Although the function remains rather broad, we can hypothesize that this gene is involved in the amide metabolic process. Finally, the RIIAD1 (ENSG00000178796) gene appears in the modules ALL_331 and COM_380. The top 5 enriched terms in ALL_331 are sperm capacitation, sperm motility, flagellated sperm motility, cilium movement involved in cell motility and cilium or flagellum-dependent cell motility. Five out of nine genes are annotated with a cellular component of the motile cilium. We can hypothesize that RIIAD1 is involved in the sperm motility. In fact, a recent paper mentioned RIIAD1 as co-expressed with the a-kinase anchor protein 3AKAP3, a gene which knockdown was shown to induce infertility of male mice [55, 56]. A visualization of the top 30 enriched terms and an overview of GO terms is available in Additional file 2 and Additional file 6 for all modules we studied in detail.

**Figure 4:**
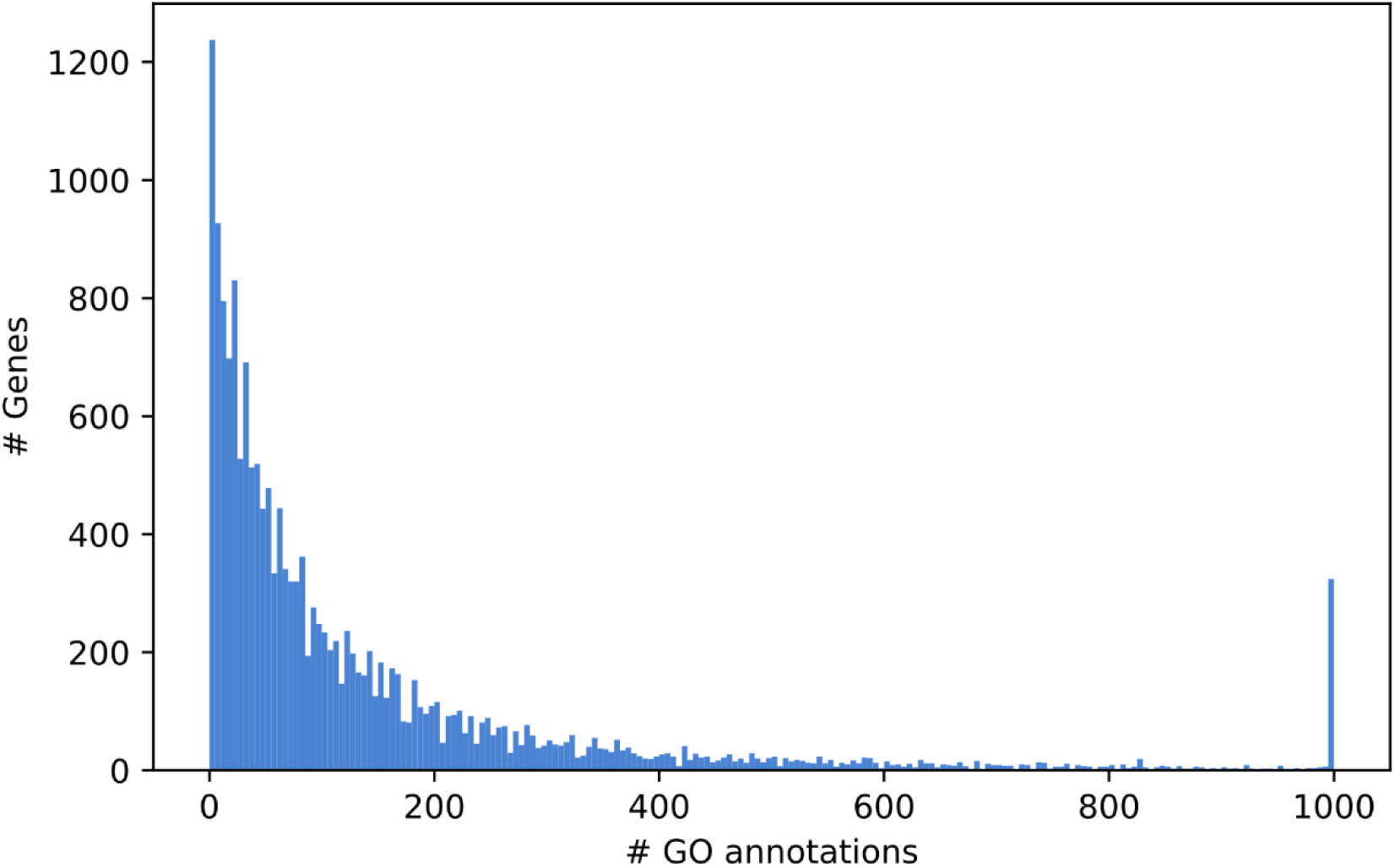
Distribution of GO annotations for all genes included in the composite *H. sapiens* network. While the majority of genes is well-annotated, 345, 245, 181, 210, 256 and 167 genes have zero, one, two three, four and five GO terms, respectively. A total of 319 genes have more than 1000 GO-annotations and were stacked at the last bin. The histogram was generated with a bin size of 5.

**Figure 5:**
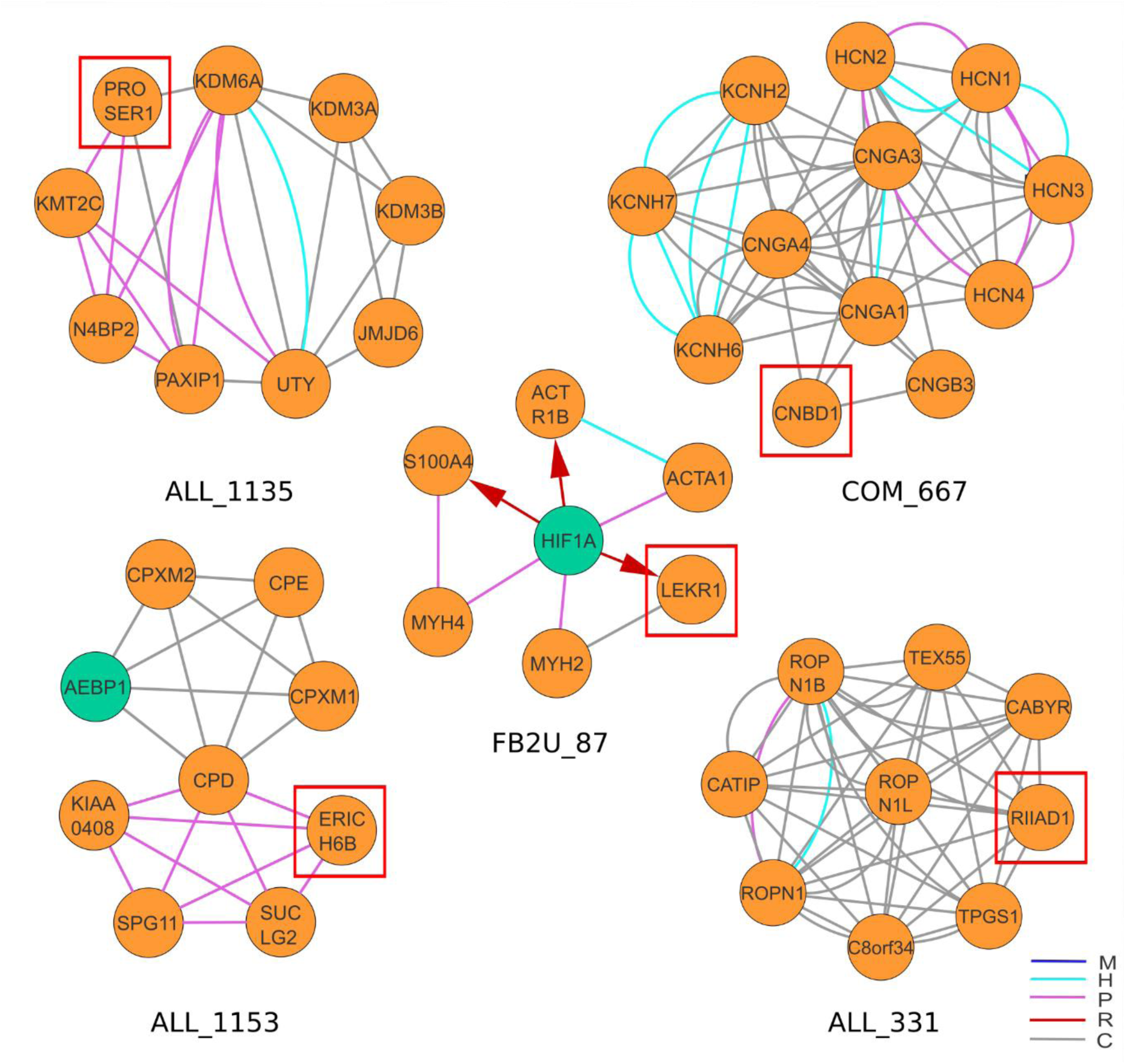
Functional prediction for poorly functionally characterized genes based on their module context. Based on the guilt-by-association principle, we created hypotheses on the biological function of these genes based on their module context. We derived functional predictions for PROSER1, CNBD1, LEKR1, ERICH6B and RIIAD1.

### Dynamic modules associated with hypoxia in three cancer cell lines

Hypoxia can lead to a variety of different responses in which cells can develop tolerance to severe tissue damage and might in turn promote aggressive cancer phenotypes [57, 58]. Such damaged cells can be embedded in the tumor microenvironment and influence the treatment effectiveness [58, 59]. To contextualize the results obtained with SUBATOMIC, we choose a study that investigated the influence of cycling and chronic hypoxia on gene expression in melanoma (WM793B), ovarian cancer (SK-OV-3), and prostate cancer cell lines (PC-3) [58]. Chronic hypoxia is characterized by a permanent oxygen depletion and was modeled in the study with an permanent ambient oxygen concentration of 1%. In cyclic hypoxia, the availability of oxygen varied between 1% and 21% oxygen with a switch at six different time points. A permanent oxygen concentration of 21% was used as control condition. We chose two main prioritization schemes to select modules potentially involved in hypoxia: one based on enriched GO terms and the other based on expression data.

We first filtered all modules based on at least one enrichment for a GO term containing the ‘hypoxia’ key word (Additional file 3). Hence, we identified 78 modules, further referenced as the ‘hypoxia GO set’. We then investigated how tightly connected this set of modules was in the superview analysis (Table 2). We used the number of interactions between modules in our selected set and compared it against a background of 1000 randomly selected sets of modules of the same set size. This allowed us to see whether modules in the ‘hypoxia GO set’ are more connected than expected by chance. We calculated the upper boundary of a 95% confidence interval on the mean and standard deviation of the random set and a fold change comparing the interactions from the ‘hypoxia GO set’ with this upper boundary. We observed that modules of the ‘hypoxia GO set’ have about 18 times more edges between pairs of modules inside this set than random. Especially regulatory edges such as TF-target interactions and miRNA-mRNA interactions between these modules are respectively 21 and 19 times more often observed than expected by chance. This revealed a strong connection between those modules, indicating complex regulatory mechanisms behind hypoxia. Next, we investigated whether the ‘hypoxia GO set’ compared to other modules is enriched for significantly differentially expressed (DE) genes in chronic hypoxia with a minimum fold change greater than two. We did not apply this to cyclic hypoxia because the number of DE genes was quite low. A hypergeometric test was used comparing DE genes appearing inside the ‘hypoxia GO set’ with DE genes appearing in all other modules. We found a significant overexpression of DE genes in all the cell lines: WB793B (fold change 16.99, p-value 3.54E-38), PC3 (fold change 9.58, p-value 1.47E-36) and SK-OV-3 (fold change 7.74, p-value 1.43E-30). These results highlighted that with a combination of superview analysis and functional annotation, we can already filter for condition-specific modules even without expression data integration.

**Table 2:** Modules in the ‘hypoxia GO set’ show higher interconnectivity than expected by chance.

| Edge type | Hypoxia GO set | Random mean | Random std | Confidence down | Confidence up | Fold Change |
| --- | --- | --- | --- | --- | --- | --- |
| M-type | 490 | 22 | 24 | 21 | 24 | 21 |
| R-type | 27476 | 1425 | 780 | 1377 | 1474 | 19 |
| H-type | 428 | 17 | 85 | 12 | 23 | 19 |
| C-type | 2772 | 202 | 216 | 189 | 216 | 13 |
| P-type | 872 | 52 | 70 | 48 | 57 | 15 |
| Total | 32038 | 1720 | 880 | 1665 | 1774 | 18 |

We calculated mean, standard deviation and a 95% confidence interval based on a distribution of 1000 randomly sampled sets of modules. Confidence up = upper boundary of the confidence interval, Confidence down = lower boundary of the confidence interval, Fold change = comparison of the number of superview interactions in the 'hypoxia GO set' with the upper boundary of the random distribution. We observed a high fold change for all edge types in the 'hypoxia GO set'.

Subsequently, we took a closer look at three selected modules in the ‘hypoxia GO set’ (Figure 6). For example, the module COM_256 resembles a co-functional protein complex that has 68 enriched GO terms including the most enriched hydroxylysine metabolic process, peptidyl-proline 4-dioxygenase activity as well as L-ascorbic acid binding (see supplement for full list of enriched GO terms). Five out of twelve genes are involved in response to hypoxia as well as response to decreased oxygen levels. The module is mostly regulated by three TFs: androgen receptor (AR), hypoxia inducible factor 1 subunit alpha (HIF1A) and endothelial PAS domain protein 1 (EPAS1), also known as hypoxia-inducible factor 2-alpha. HIF1A and EPAS1 are known to facilitate cellular adaptation to hypoxia and regulate many hypoxia related genes in a variety of tissues [60–64]. Also AR was shown to act as ligand-dependent TF that confers to resistance against AR-targeted cancer therapies under hypoxic conditions [65, 66]. Many module genes were differentially expressed, namely the procollagen-lysine,2-oxoglutarate 5- dioxygenase 1 and 2 (PLOD1, PLOD2), the prolyl hydroxylase (EGLN3), as well as prolyl 4- hydroxylase subunits alpha 1 and 2 (P4HA1) and (P4HA2). PLOD1 and PLOD2 were shown to be involved in hypoxia-induced metastasis and glioblastoma tumor progression [63, 67]. EGLN3, which was upregulated in the chronic state in the SK-OV-3 cell line, catalyzes oxygen-dependent hydroxylation of the hypoxia induced factor (HIF) [68, 69]. Other genes such as the prolyl 3-hydroxylases, P4HA1 and P4HA2 were known to hydroxylate the 564-proline residue in the α-subunit of HIF [70]. Hence, concluded that COM_256 is strongly involved in the reaction to hypoxia and shows coherent but slightly different expression across the three cell lines.

**Figure 6:**
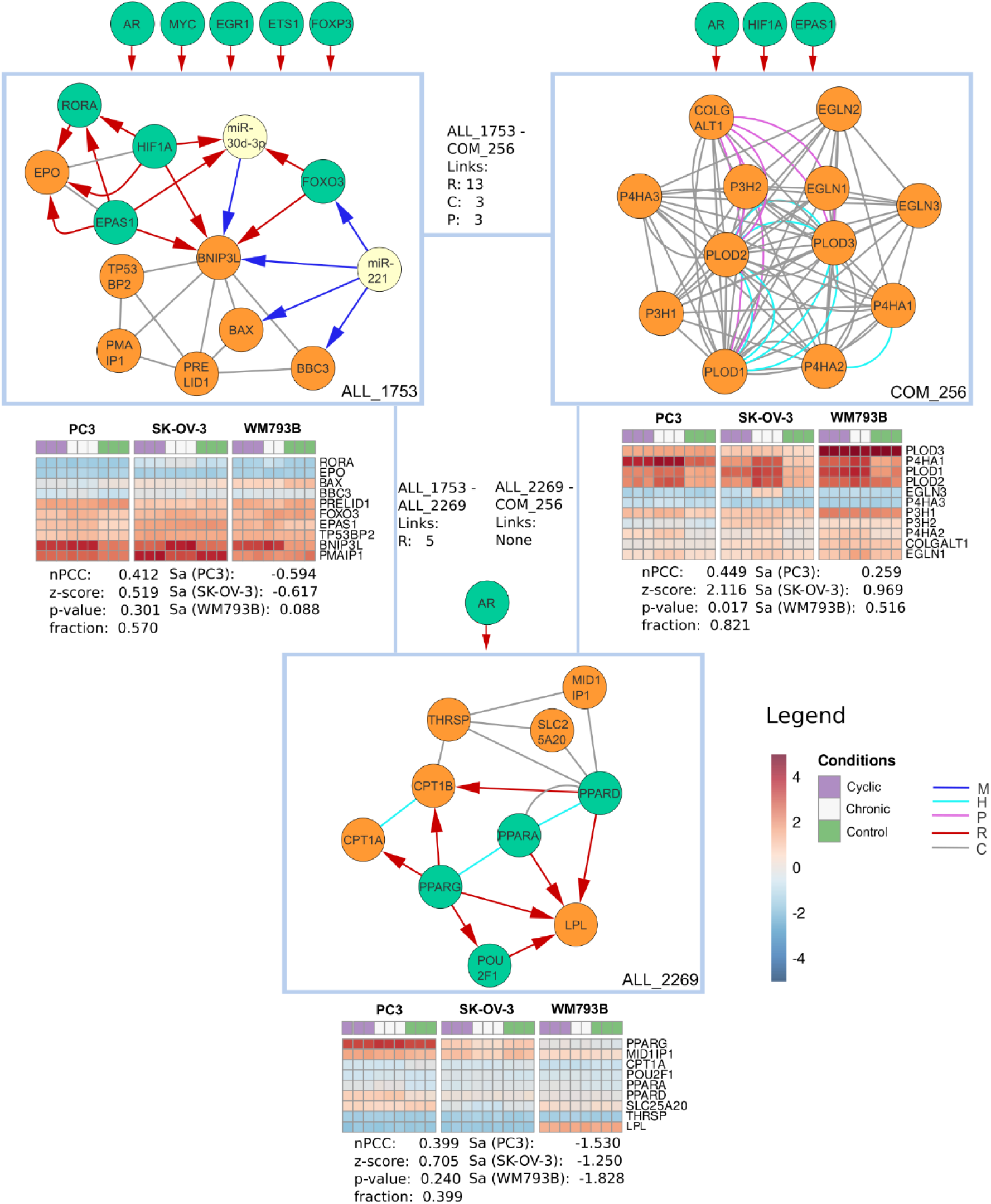
Visualization of three superview connected modules from the ‘hypoxia GO set’. ‘Links’ indicate how many superview interactions exist per type between two modules. Regulators targeting at least 5 genes that were not present in a specific module are shown outside the boxes. nPCC indicates the correlation of genes within a module in all hypoxia samples together with a z-score derived by comparing this value with random modules of the same size as well as an p-value and the fraction of edges in a module for which expression data was available for both genes. The activity score Sa was displayed for each of the three cell lines PC3, SK-OV-3 and WM793B. Expression is shown for all module genes for which expression values were available.

We then investigated the ALL_1753 module. The module contained 177 significant GO term enrichments including cellular response to hypoxia and cellular response to decreased oxygen levels for six genes. It is centered around the BCL2 Interacting Protein 3 Like gene (BNIP3L), which is differentially expressed and targeted by HIF1A and EPAS1. The module contained three interesting feed-forward-loops, where HIF1A, EPAS and the forkhead box O3a FOXO3 target BNIPL3 and miR-30d-3p that in turn also regulates BNIP3L. Moreover, miR-30d-3p is known to be involved in hypoxia and directly regulate AR [71]. FOXO3 is activated in response to hypoxic stress (Bakker et al., 2007). Another DE gene in this module is the retinoic acid receptor-related orphan receptor (RORA), regulated by HIF1A and EPAS1. RORA is known to be induced by HIF1A and it plays a role in the nuclear accumulation of HIF1A [74]. Finally, miR-221 regulated FOXO3, BNIPL3, the bcl-2-binding component 3 (BBC3) and the apoptosis regulator BAX and exerts cytoprotective effects in hypoxia-reoxygenation injury [75]. Hence, we concluded a strong involvement of ALL_1753 in response to hypoxia and that the hypoxia response of BNIPL3 might be driven by the involvement of three regulatory feed-forward loops.

At last, we investigated the ALL_2269 module. The module was enriched for 85 GO terms including the most enriched terms carnitine shuttle and carnitine O-palmitoyltransferase activity for two and three genes as well as positive regulation of fatty acid metabolic process that involved half of the module genes. The response to hypoxia and decreased oxygen levels was enriched due to the presence of three module genes. The module is centered by three homologous peroxisome proliferator-activated receptors, PPARG, PPARA and PPARD. Especially PPARG was shown to be activated under hypoxic conditions in correlation with HIF1A in lung cancer and hepatocellular carcinoma [76, 77]. It regulated the differentially expressed gene carnitine palmitoyltransferase 1A (CPT1A) and its homolog CPT1B, shown to regulate prostate cancer growth under hypoxic conditions [78]. While this module does not show strong dysregulation in the three analyzed cancer types, it contained genes and interactions highly relevant in reaction to hypoxia as demonstrated in other studies.

All three modules were connected by many edges in the superview and share a similar set of regulators. We demonstrated that the modules found by our GO term based approach are highly relevant in the hypoxia context which is supported by the increased amount of hypoxia specific DE genes. In the modules we identified regulatory structures such as feed-forward loops involving interactions from complementary omics layers that help to explain and interpret the observed expression and allow to generate mechanistic hypotheses for hypoxia induced mechanisms.

In a second prioritization approach, we wanted to use the dynamic response of genes towards a stimulus or condition as selection criteria (Additional file 4). We implemented a ‘module activity’ *S_a_* approach that can capture the response of modules to a changing condition, for example based on differential expression data between two conditions [79] (see also methods). To find a set of highly hypoxia related modules, we filtered for modules with a positive activity score *S_a_* in all three cell lines. This resulted in a set of 52 modules that we further refer to as the ‘hypoxia activity set’. Next, we analyzed the superview connections within the ALL modules in the same way as for the ‘hypoxia GO set’ (Table 3). We observed a 28 times higher connectivity between the activity modules as compared to random sets of the same size. While the number of interactions for M-type and R-type interactions was similar to the ‘hypoxia GO set’, the undirected interaction types H, C and P were much more enriched with an 90, 69 and 88 times higher number of connections, respectively. Furthermore, the ‘hypoxia GO set’, and the ‘hypoxia activity set’ have 12 modules in common.

**Table 3:** ‘Modules in the ‘hypoxia activity set’ show higher interconnectivity than expected by chance.

| Edge type | Hypoxia | Random mean | Random std | Confidence down | Confidence up | Fold Change |
| --- | --- | --- | --- | --- | --- | --- |
| M-type | 134 | 7 | 9 | 6 | 7 | 18 |
| R-type | 8494 | 458 | 330 | 437 | 478 | 18 |
| H-type | 624 | 5 | 25 | 4 | 7 | 90 |
| C-type | 5008 | 66 | 112 | 59 | 73 | 69 |
| P-type | 1676 | 17 | 35 | 15 | 19 | 88 |
| Total | 15936 | 552 | 370 | 529 | 575 | 28 |

We calculated mean, standard deviation and a 95% confidence interval based on a distribution of 1000 randomly sampled sets of modules. Confidence up = upper boundary of the confidence interval, Confidence down = lower boundary of the confidence interval, Fold change = comparison of the number of superview interactions in the 'hypoxia GO set' with the upper boundary of the random distribution. We observed a high fold change for all edge types in the 'hypoxia GO set'.

We then inspected three modules identified in the activity hypoxia set with regard to their relevance for hypoxia (Figure 7).

**Figure 7:**
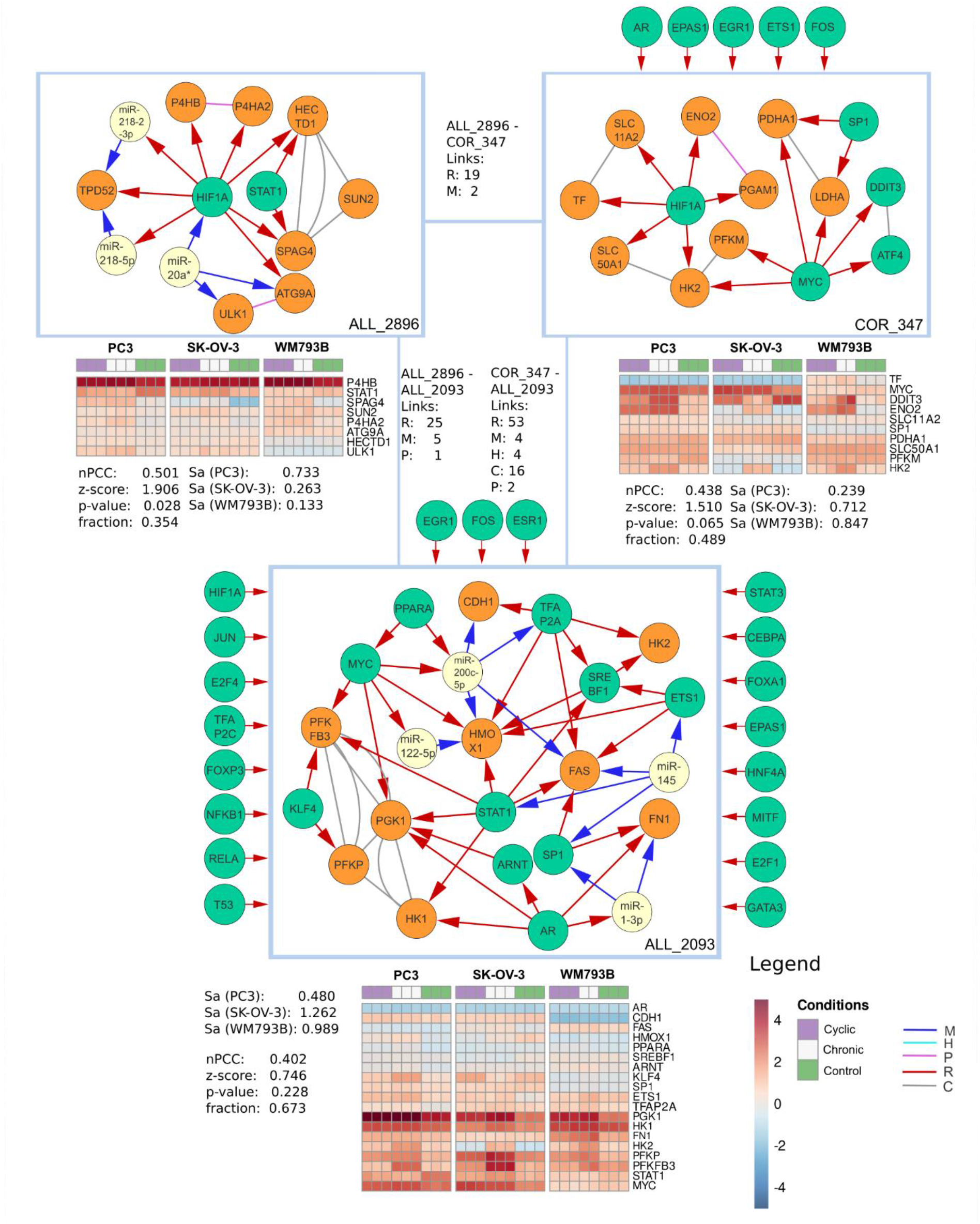
Visualization of three superview connected modules from the ‘hypoxia activity set’. ‘Links’ indicate how many superview interactions exist per type between two modules. Regulators targeting at least 5 genes that were not clustered in a specific module are shown at its top. nPCC indicates the correlation of genes within a module in all hypoxia samples together with a z-score derived by comparing this value with random modules of the same size as well as an p-value and the fraction of edges in a module for which expression data was available for both genes. The activity score Sa was displayed for each of the three cell lines PC3, SK-OV-3 and WM793B. Expression is shown for all module genes for which expression values were available.

The COR_347 module is dominated by HIF1A and MYC, each regulating 6 targets. The module contained 159 significant GO term enrichments including cellular response to hypoxia and cellular response to decreased oxygen levels for four and three genes, respectively. Besides HIF1A and the MYC proto-oncogene (MYC), the module is regulated by the *fos* proto-oncogene (FOS) with six regulatory interactions. MYC is one of the master regulators targeting 46% of all modules. It plays an important role in the development of cancer and regulates members of the hypoxia inducing factor protein family [80]. The hexokinase 2 gene (HK2) is regulated by MYC and HIF1A and shows dysregulation in the expression data. It was recently shown to be an important target of HIF1A in an oxygen-reduced environment [81].

ALL_2896 mostly contained COR and FFL subgraphs dominated by HIF1A. It was enriched for 11 GO annotations including procollagen-proline 4-dioxygenase activity and peptidyl-proline 4-dioxygenase activity. HIF1A regulated the differentially expressed gene P4HA1, which also occurred in COM_256 in the ‘hypoxia GO set’ in a co-regulatory manner with the prolyl 4-hydroxylase beta polypeptide gene (P4HB). Another regulator in this module was the signal transducer and activator of transcription 1 gene (STAT1). It regulated the sperm associated antigen 4 (SPAG4) together with HIF1A. While SPAG4 was not differentially expressed with a fold change greater than two, we still observed a consistent reduction in expression across all three cell lines.

ALL_2093 is enriched for 331 GO terms. The most enriched terms included regulation of metanephric cap mesenchymal cell proliferation and negative regulation of leukocyte adhesion to arterial endothelial cell, however the most enriched 15 terms only contain one gene. The response to hypoxia as well as to decreased oxygen levels included six genes. The module mostly consisted of COR and FFL subgraphs. Many of its genes connected to hypoxia already appeared in the modules described above, such as AR, MYC, HK2, STAT1 and PPARA. Another interesting gene is the krüppel-like factor 4 (KLF4). It was differentially expressed together with its target 6-Phosphofructo-2-Kinase/Fructose-2,6-Biphosphatase 3 (PFKFB3), which in turn is also regulated by STAT and MYC. KLF4 was shown to be involved in Hypoxia- induced vascular smooth muscle cells migration [82]. However, while PFKFB3 is differentially expressed in PC-3 and SK-OV-3 cell lines, KLF4 is only significantly overexpressed in PC-3. Besides regulation via STAT1 and MYC, the PFKFB3 promotor also contains HIF1A binding sites and might not be dependent on KLF4 as activator to be overexpressed under hypoxic conditions [83]. Furthermore, fibronectin 1 (FN1) is overexpressed and regulated by two FFLs including AR, SP1 and miR-1-3p. The latter was show to be downregulated under hypoxic conditions in mouse lung tissues [84]. The specificity protein 1 (SP1) was described to also be regulated by HIF1A directly and is required for hypoxia induced transcription of other downstream genes [85].

## Discussion

We developed SUBATOMIC, an integration pipeline that decomposes multi-omics networks into topological modules and their interactions using composite subgraph clustering and statistical and functional analysis. The obtained modules are further embedded within their network and regulator context in a superview analysis as well as functionally annotated and visualized. In a post-processing step, we contextualized the obtained modules with condition-specific data in three hypoxia cell lines and calculated activity as well as expression correlation scores. Compared to our previous integration framework, SUBATOMIC contains many improvements [28]. Most importantly, we automated the workflow and integrated it into a general Snakemake pipeline. While the previous version was generated for one use case only, we now can perform the analysis workflow from decomposing a composite network up to the analysis of modules in one single execution. Moreover, it is applicable to multi-edge networks of any species. We adapted the superview analysis to output summary of connections between modules and regulators. We also improved upon the functional characterization of modules by automatizing GO term enrichment. Since run-time can be an issue on large interaction networks, we parallelized time critical steps in the pipeline and improved the scalability and computability. Moreover, we added scripts to support the visualization of modules in Cytoscape. To calculate module activity, we developed a post-processing step that calculates a biological activity score.

We applied SUBATOMIC to a composite network consisting of human TF-target gene, miRNA-mRNA, protein-protein, co-functional and homologous interactions. Our approach yielded 5586 modules. The majority of modules was enriched for GO terms, demonstrating that module topology and biological function are closely interrelated, since the clustering was performed on topological features. Most modules obtained were COM modules, made up of undirected edges. This is expected because we included more undirected than directed interactions. Moreover, we observed that in most of the cases the interactions in a three-node subgraph came more often from the same input network rather than from different networks. This is due to the fact that the input networks do not possess exactly the same set of nodes and can also have a different number of interactions. Instead of using the full HumanNet database, we included only high-quality interactions. This balances the number of interactions to a comparable amount by setting cut-offs on the quality values.

Given the modules and their functional annotation, we demonstrated how SUBATOMIC can be used to predict the function of unannotated gene. Out of 53 weakly annotated genes that also appeared in the inferred modules, we selected PROSER1, CNBD1, LEK1, RIIAD1 and ERICH6B for an in-depth analysis. While some of our predictions such as the chromatin modification role of PROSER1 are very novel, others were supported by recent publications such as the potential involvement of RIIAD1 in sperm motility. However, the functional characterization based on guild-by-association principle generates guiding functional hypotheses that still need experimental conformation.

In the case of contextualized modules, we put forward two prioritization strategies: either through enrichment of related GO terms or the module activity score. We showed that modules that share GO terms such as hypoxia are strongly interconnected and accumulate DE genes for related conditions. Also genes not annotated with the specific GO terms but relevant for the condition can be detected due to close distances inside one module. However, when genes are not well characterized or do lack specific GO key words linking them to a condition, a GO term based prioritization strategy might miss out on important modules. We further developed a more data driven prioritization scheme by implementing module activity based on differential expression data. Using the activity score, we found a small set of modules relevant in the hypoxia context. This set was strongly interconnected and partially overlapped with the GO term based set of genes. While modules on itself show a static view and give insights on what is possible in an organism, contextualizing adds condition specific dynamicity. For example, despite the fact that one of the main driving hypoxia genes HIF1A was not present in the most current probe annotation for the expression array from the hypoxia study, it was detected in many modules based on GO term annotation and activity. Especially genes targeted by HIF1A revealed high differential expression in HIF1A containing modules. Our results indicate that we can prioritize the large amount of modules in different ways to end up with sets of modules highly relevant for a condition or disease context. With the activity prioritization method, we were able to find modules with different topology strongly connected in the superview. We showed that many genes in these modules are already known to play important roles in hypoxia. Moreover, contextualizing the modules with expression data from three different cell lines revealed that the activation of response mechanisms can differ and that different parts of a module can be active in different tissues. For example, EGLN3 is a known hypoxia induced factor and showed dysregulation only in the SK-OV-3 cell line (see COM_256). In ALL_2093, KLF4 is only weakly expressed in WM793B while its target PFKB3 is strongly expressed in all three cell lines under hypoxic conditions, thus other regulators such as MYC and STAT also targeting PFKB3 might have a stronger regulatory influence. Overall, the combination of annotated SUBATOMIC modules based on different topologies, their superview connections, their contextualization with expression data and their visualization among different conditions and cell lines delivers a versatile tool to deeply investigate multi-edge networks.

SUBTATOMIC is not limited to the data types described in this study and offers additional analysis opportunities. While we restricted our analysis to genes and miRNAs, it is possible to add any type of nodes and interactions such as metabolite interactions, lncRNA interactions, or siRNA interactions as long as the input networks share a common set of nodes for intersection. Moreover, while we used a static prior network that was contextualized in a later step to add dynamicity, it is possible to directly analyze a dynamic composite network by including condition- or patient specific association networks inferred from context-specific high-throughput data such as transcriptomics or proteomics, e.g. co-expression networks at either bulk or single cell level. Also hybrid approaches are possible combining public databases for some interaction types with condition specific interactions for others.

Broadly, two main distinctions can be made when it comes to module inference: on the one hand, methods generate modules directly from experimental read-outs such as expression data or use prior networks as base for the inference. On the other hand, the clustering can be based on only one data modality or include multiple ones. Methods such as WGCNA, lmQCM and TPSC are examples for methods that generate expression based co-expression clusters [38, 86, 87]. While they are widely used and are shown to produce modules that correlate to many biological features, these modules are often very large, do not consider causal regulatory interactions or have no multi-omics integration. Other methods go one step further and additionally integrate protein-protein interactions [41, 88]. They combine the clustering on expression data with protein-protein interaction networks to derive modules that represent merged, overlapping and independent communities [88]. While the classification of modules in independent, overlapping and merging communities already give some network context to the clusters, it is not as flexible with the input data. Another method used a manel test to integrate co-transcriptional regulation and protein-protein interaction networks [41]. While they demonstrated functional coherence between modules based on prior interactions, they did not consider directionality of interactions and the clustering yielded only a small number of modules. Another class of methods tries to use static prior networks as base and uses expression data in a contextualization step to find active subnetworks [89, 89–91]. Some more of these methods are reviewed by other authors [92]. Compared to existing methods, SUBATOMIC tries to address open issues and creates an comprehensive analysis framework covering various aspects of module inference. It is based on a topological clustering approach that allows to interpret clausal relationships between gene and emphasizes different regulatory mechanisms. It introduces flexibility allowing for operation on literature-defined prior networks as well as on expression-derived association networks. Moreover, It can include all types of nodes and biological interactions in an integrated manner. Networks are divided into a large number of small and interpretable modules with district topological properties, while still keeping track of their global network context and regulators. Furthermore, static network modules can be inferred and contextualized with expression data using ECD, nPCC and activity scores. This combination makes it a unique and outstanding method in the field of composite network clustering.

While SUBATOMIC was shown to be able to answer many biological questions, it also comes with some limitations. Input prior networks are often incomplete and might complicate some interpretations. If a regulator is module specific but also relevant for a function not represented in the input networks, it might be less specific in more complete networks. However, we expect that interaction databases continuously grow, and thus multi-edge networks will become more and more complete. Furthermore, if networks do not overlap for a certain quantity of nodes, the derived subgraphs will mostly assemble in modules from separated interaction types. Also this issue will be solved with a growing amount of databases. Another limit is computability. We used SCHype for our clustering algorithm and demonstrated that it was able to process more than 750000 subgraphs in an adequate amount of time. However, the number of detected subgraphs grows super-linear with larger networks. Thus, there exists an upper limit in network size that is still computable. Furthermore, while we parallelized GOATOOLS to annotate several modules at a time, the gain of computational speed was accompanied by an increase in space consumption. This limits the number of cores that can be used for parallelization. This will be addressed in a future version integrating a more space-efficient annotation tool.

## Conclusion

In conclusion, we developed an automated subgraph clustering framework that takes the basic building blocks of interactions and clusters them into modules. The modules are further characterized and contextualized by superview calculation, regulator analysis, GO term enrichment and module activity scoring. SUBATOMIC can be used to investigate conditions and diseases, find interactions between functionally related modules and derive novel gene functions for uncharacterized genes. The main limiting factor is the availability of interconnected networks. We believe this issue will be solved in time with an ever-increasing number of interactions being discovered. Our approach distinguishes itself from other module inference methods based on clustering expression data or network edge rations, by clustering topological building blocks to create a high number of small and easily interpretable modules with different regulatory properties, while still keeping the overall network connections in mind.

## Methods

The main analysis workflow has been integrated into one Snakemake pipeline. A Snakemake workflow diagram is available in the Additional file 6, and a schematic overview can be found in Figure 1. All software, including some scripts for pre- and post-processing analysis, as well as a Docker version are made available on GitHub (https://github.com/CBIGR/SUBATOMIC).

### Subgraph detection

For subgraph detection, we used the ‘Index based subgraph algorithm’ (ISMAGS) [44, 93]. We followed the subgraph representation used in ISMAGS, where a three-node subgraph is represented as a three letter code, which specifies that a given edge originates from a certain set of input interactions. All interactions of one input network need to be either all directed or all undirected (e.g. all interactions in a TF-target network should be directed, all interactions in a protein-protein network should be un-directed). Each input network is characterized by a unique letter representation that can freely be chosen. As an example, the subgraph RRP would contain one directed edge from network R, another directed edge from network R as well as an undirected edge from the protein-protein interaction network. A letter for a directed network can be set to lower case to indicate that the direction is reversed (see ISMAGS paper). Due to symmetry, some subgraphs are redundant to one another (e.g., PPC, PCP and CPP represent the same subgraph). A custom-made script calculated a non-redundant set of subgraph representations based on a provided list of directed and undirected network letters. This set is then used by the pipeline as guide to search for subgraphs and can be further fine-tuned by the user to remove additional unwanted subgraphs. The three-node subgraphs are then identified by ISMAGS [44]. ISMAGS takes for each iteration the three-letter subgraph representation and the parts of the composite network that contain interactions for this subgraph. It outputs all three-node subgraphs that satisfy the defined representation. Besides three-node subgraphs, we also searched for two-node subgraphs that possess special properties: i.e. all pairs of nodes where each node contains a directed edge pointing at the other node (DD type) and all pair of nodes connected by one undirected and one directed edge (DU type).

### Subgraph clustering and module inference

The subgraphs produced by ISMAGS were subsequently grouped into one of the following subgraph types: complex subgraphs (COM), feed forward loop (FFL), co-pointing subgraphs (COP), co-regulated subgraphs (COR), circular feedback subgraph (CIR), feedback undirected subgraph (FBU) and feedback 2 undirected subgraph (FB2U) and two-node feedback subgraph (2FB) (Figure 1, [28]). For example, the COM type contains all subgraphs that exclusively consist of undirected edges. Given the undirected network letters C, P and H, any combination of these three letters that lead to a set of non-redundant subgraphs was grouped together into the COM type. Each module type consists of specific subgraphs types and serve as the input for the following clustering.

We inferred clusters for each of the above defined subgraph types separately as well as on the union of all classes (ALL) using the SCHype algorithm [45]. This algorithm is based on the Perron-Frobenius theorem and clusters a hypergraph solving an optimization problem by maximizing the edge-to-node ratio in each cluster for a network [94]. The input is a hypergraph were each hypernode represents a three-node subgraph calculated by ISMAGS or a two-node subgraph. SCHype was run with default settings (P=1) and output several modules for each of the eight classes of modules and for clustering all subgraphs together. These modules were further filtered: clusters containing 5-50 genes were kept and modules containing more than 90% of homologous edges were excluded for subsequent analyses.

### Superview calculation

The superview step characterizes interactions between modules as well as between regulators and modules. Each module was compared to every other module by counting how many edges per input network were shared between two modules. This value was compared against the shared edge count in a random sampling. In the sampling we generated 1000 times two random modules, both having an equivalent amount of nodes as the two modules under investigation. The derived distribution was used for a z-score transformation with 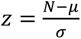 given the mean *μ* and the standard deviation σ from the random distribution. The *z*-score was further evaluated by calculating a p-value (significance cut-off: 0.05) for that distribution with a right tailed test 1 – *CDF*(*z*) where *CDF* is the cumulative distribution function. The output was composed of one file per module type (ALL, COM, COR, …) containing interactions between every module of this type and all other modules for each input network and displayed the count of shared interactions, z-score, and p-value. If no interactions exist between two modules, the z-score was set to 0 and the p-value was set to 1.

The superview calculated three more outputs that characterized the relationship between modules and regulators. For each regulator (TF or miRNA), we calculated the RF-module connection strength for each RF-module pair and each interaction type with 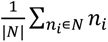 with N being all module genes and *n_i_* = 1 if an edge exist, else *x* = 0. This gave a fraction on how many interactions between a TF and a module existed and was used to find regulators that are strongly connected to a module or a set of modules. Another analysis displayed the fraction how many distinct TF or miRNA target one certain module for each module with 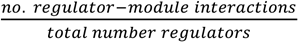. This gave a module-specificity and allowed to investigate whether a module was targeted by a few or many regulators. Another analysis displayed how many modules are targeted by a certain regulator and shows a fraction how this compared to the total number of derived modules. For each regulator, we calculated the regulator specificity by 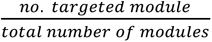, which allowed to investigate which TF and miRNAs targeted a wide range of modules as opposed to some that were specific for a single module or a small number of modules.

### Functional enrichment analysis

For each module, SUBATOMIC performed a functional enrichment to gain insights into its biological relevance. We used the Python implementation of GOATOOLS to calculate the enrichment of GO terms for each module [46]. We provided three options as enrichment background: all genes present in one specific type of modules, all genes present in the input networks or all genes written in a user-specified file. For our analyses we used a user-defined input containing all annotated human genes as input. Results were summarized in one file per module. Only results with a corrected p-value > 0.05 were kept based on Benjamini and Hochberg FDR [95]. Additionally to the standard GOATOOLS output, we provided a rank for the ascendingly sorted p-values for each module, since p-values can strongly differ between modules and enriched functionality depends on which processes are well annotated as well as how many GO terms are available. The rank allowed to filter for the top n entries per module. We further reported the log2 fold change for each significant GO term.

### Visualization

To visualize the modules in Cytoscape, we provide a number of files that can be imported. The most important file is an nnf file containing the network representation of the modules. Additionally, we generated a noa file to annotate each node with its type (TF, gene or miRNA), its gene name and a short optional functional description. Each run of the pipeline also resulted in a Cytoscape style sheet in xml format that can be imported as well to format the network in a way consistent with the provided information in nnf and noa file. The xml file can also be adapted for more customized style choices.

### Run time considerations

Operations on graphs often come with a high computational cost. Several steps in the pipeline were parallelized, but some bottlenecks remain. The subgraph detection algorithm is highly efficient and can find millions of subgraphs in less than a minute. SCHype can cluster hundreds of thousands of subgraphs, but its runtime increases super-linear Furthermore, the superview calculation comparing all modules against each other and the functional annotation are the most time-consuming steps. Since these steps process one module at a time, we parallelized them in a way that each module can be processed by a different core. In principle, as many modules can be processed in parallel as there are cores available. However, since each separate process needs a certain amount of memory, a careful balance between the number of cores available and the amount of memory must be taken in order to find a suited number of cores for parallelization. For our application on a *H. sapiens* composite network, we run SUBATOMIC with eight cores and 70GB RAM on a 2 x 18-core Intel Xeon Gold 6240 (Cascade Lake @ 2.6 GHz) processor, which resulted in two days of runtime.

Construction of the composite *H. sapiens* network

We unified five different types of interactions from different sources into one composite network representation (Table 1). From OmniPath, we included 53232 TF-target gene interactions formed by 526 regulators and 14488 target genes (access 17.01.2022) [19, 20]. We included all interactions in the evidence classes A, B and C from DoRothEA as well as the TF-target gene and TF-miRNA interactions. From the same database, we included 11085 miRNA-target interactions between 850 miRNAs and 3210 target genes (access 17.01.2022). The homology between genes was retrieved from Ensembl based on the GeneTree pipeline [49]. This pipeline takes a reciprocal best BLAST approach in a simple case but also considers more complex ontologies by resolving one-to-many and many-to-many relations. We applied a minimum reciprocal sequence identity of 50% as threshold to include homologous interactions between two genes. We only considered genes involved in at least one of the other data sets included in this analysis. Moreover, we added homologous miRNAs with identical seed sequence. This summed up to a total of 10847 interactions for 4862 genes or miRNAs. We obtained protein- protein interactions and co-functional interactions from HumanNet v2 [21, 50]. We used the log-likely-hood score (LLS) provided by HumanNet to filter for interactions with LLS <= 3.0. Since the number of interactions in HumanNet was magnitudes bigger than the number of regulatory interactions, this filtering steps tried to balance the number of interactions without losing to many included genes. We selected 24773 protein-protein interactions formed by 10950 genes and 66373 co-functional interactions formed by 10,683 genes. For the undirected interaction sets, duplicates were merged (e.g., A-B and B-A is equivalent for undirected interactions). This removed a total of 6 protein-protein interactions, 28 co-functional interactions and 39 homologous interactions. We mapped all genes to Ensembl identifiers to make them comparable between the networks. For miRNAs, we kept the standard naming convention (e.g. hsa-miR-600e), which avoided potential overlaps with Ensembl gene identifiers. We omitted genes that could not be mapped to an Ensembl ID. The genes included in the analysis are based on the human genome version 38. We included annotated genes on chromosomes 1-22 as well as X and Y. All interactions were merged into one file to create the composite network. In this file, each interaction was represented by the two interacting nodes as well as their edge color. The edge color was represented with a network specific letter (TF-target gene interactions: R, miRNA-target gene interactions: M, homologous interactions: H, protein-protein interactions: P, Co-functional interactions: C, Additional file 1).

### The hypoxia expression data set

We used expression data from three different cell lines under cyclic and chronic hypoxia conditions to contextualize the modules (GEO: GSE53012) [58]. The Affymetrix microarray data was processed and normalized using the Single Channel Array Normalization (SCAN) [96]. It corrected the effect of technical bias, such as GC content by applying a mixture-modeling approach [96]. To calculate the biological activity, we used the p-values from the differential expression analysis from the original publication.

### Contextualization

We implemented several methods to contextualize modules with expression data from perturbation experiments or experiments with case and control samples. We calculated the average Pearson correlation coefficient nPCC between each pair of genes in a module and compared it against the nPCC of a sampling of 1000 modules to measure co-expression within a module. Next, we derived a z-score for each module nPCC and transform it into a p-value via the CDF of the standard normal distribution. We also added an implementation of the Expression Correlation Differential Score (ECD score), which highlights modules specific for an experimental condition as compared to the control condition [28]. There we subtracted for each edge in a module the Pearson correlation of case samples from the Pearson correlation of condition samples and averaged this over all edges per module. We repeated this for 1000 randomly generated modules to obtain a background distribution, which is then used to calculate a z-score that was consequently transformed into a p-values using the standard normal CDS. Given enough expression values and conditions, this allowed us to address the dynamicity of edges using the guilt by association and guilt by rewiring principles.

We implemented one additional metric compared to the previous framework to capture modules responding to a change in condition. This module activity score (*S_a_*) used p-values of a comparison between different conditions (e.g., from differential expression between case and control samples) and transformed them into a z-score *z_i_* = *θ*^−1^(1 – *p_i_*) with *θ*^−1^ beeing the inverse normal CDF [79]. For each module, we calculated the aggregate z-score *z_a_* =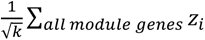 for each module with size k. For each module size, we drew 1000 random modules of the same size and used this as background distribution to calculate the normalized activity score 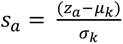. High values of *S_a_* indicated that the module can be interpreted as an active subnetwork in the specific experimental condition.

## Acknowledgements

We would like to thank Kenneth Stoop and Pieter Audenaert for their help with running the ISMAGS algorithm. Moreover, we would like to thank Hayoung Kim, Heesoo Song and Jietse Verweirder for their support in prototyping the Snakemake pipeline.

## Declarations

### Availability of data and materials

GitHub: https://github.com/CBIGR/SUBATOMIC, code of the pipeline as well as link to the Docker version

Zenodo: https://doi.org/10.5281/zenodo.6556413, raw data of the input networks and SUBATOMIC output

### Authors information

Vanessa Vermeirssen and Jens Loers have the following affiliations:

Lab for Computational Biology, Integromics and Gene Regulation (CBIGR), Cancer Research Institute Ghent (CRIG), Ghent, Belgium

Department of Biomedical Molecular Biology, Ghent University, Ghent, Belgium Department of Biomolecular Medicine, Ghent University, Ghent, Belgium

### Ethics declaration

#### Ethics approval and consent to participate

The data analyzed in this study were from public databases, so ethical approval and consent participation were not required.

#### Consent for publication

Not applicable.

### Additional information

Corresponding author Vanessa Vermeirssen

### Author’s contributions

Contributions according to the CRediT system (https://casrai.org/credit/, JL = Jens Uwe Loers, VV = Vanessa Vermeirssen):

Conceptualization: JL, VV, Data curation: JL, Formal analysis: JL, Funding acquisition: JL, VV, Investigation: JL, Methodology: JL, VV, Project administration: VV, Software: JL, Supervision: VV, Validation: JL, Visualization: JL, Writing - original draft: JL, VV, Writing – rewriting & editing JL, VV.

### Supplementary information

Additional file 1.xlsx: General description of the input networks and inferred modules

Additional file 2.xlsx: Analysis of weakly characterized genes

Additional file 3.xlsx :GO term based module analysis

Additional file 4.xlsx: Activity based module analysis

Additional file 5.xlsx: Abbreviations of gene names

Additional file 6.xlsx: Supplemental figures

## List of abbreviations

2FB: Two-node feedback subgraph
CIR: Circular feedback subgraph
COM: Complex subgraph
COP: Co-pointing subgraph
COR: Co-regulated subgraph
FB2U: Feedback 2 undirected subgraph
FBU: Feedback undirected subgraph
FFL: Feed forward loop
RF: Regulatory factor
TF: Transcription factor
COF: Co-functional interaction

The abbreviations of all gene names mentioned are listed in the supplement.

