## Additional file 6 for "SUBATOMIC: a SUbgraph BAsed mulTi-OMIcs Clustering framework to analyze integrated multi-edge networks"

**Supplemental figures**

Collection of supplemental figures. The enrichment plots were done with enrichR on an imported dataframe based on the SUBATOMIC output [1]. If there were more than 30 significant GO enrichments, only the top 30 genes were visualized.

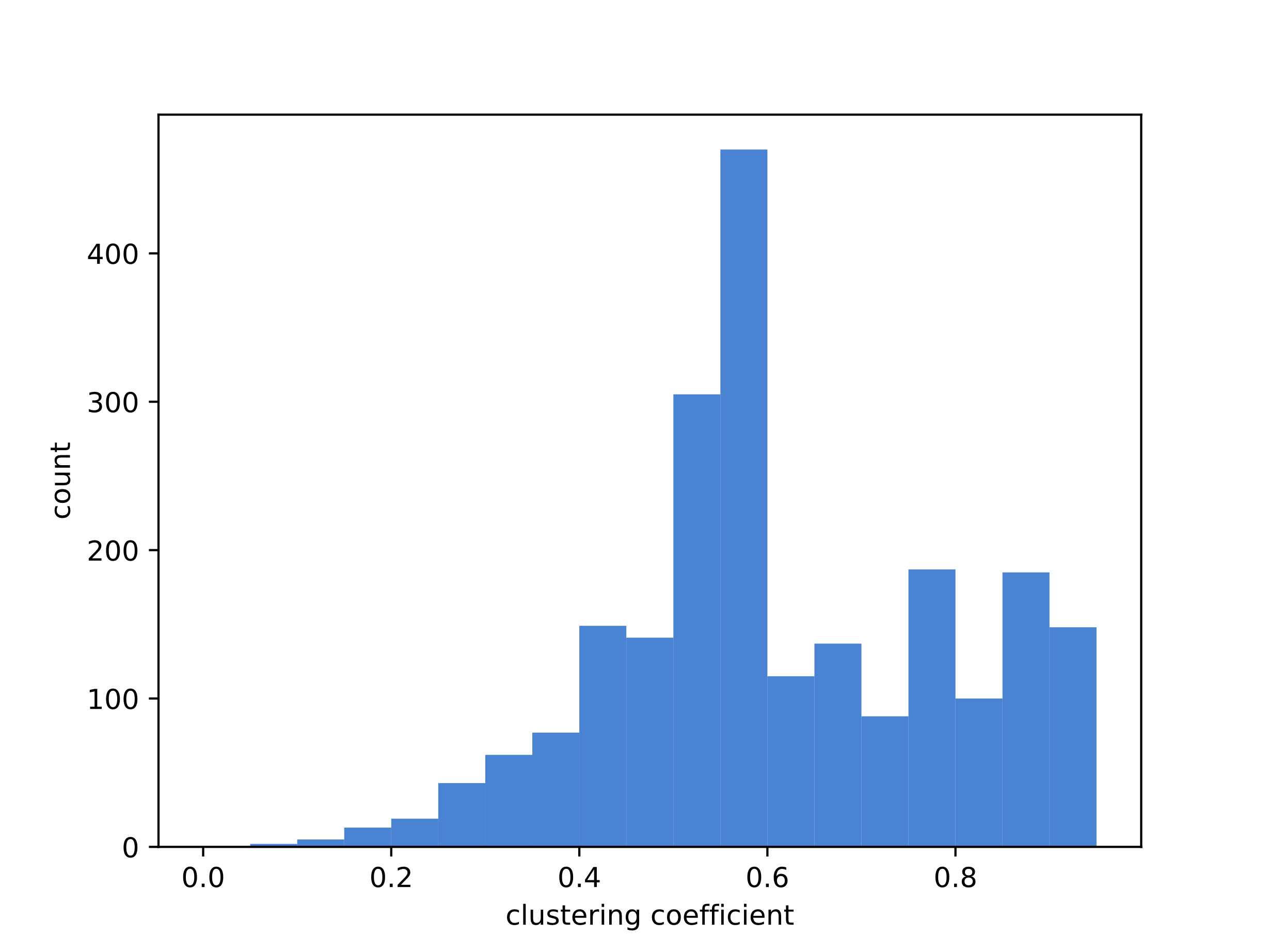

Figure 1 Distribution of the clustering coefficient for all ALL type modules.

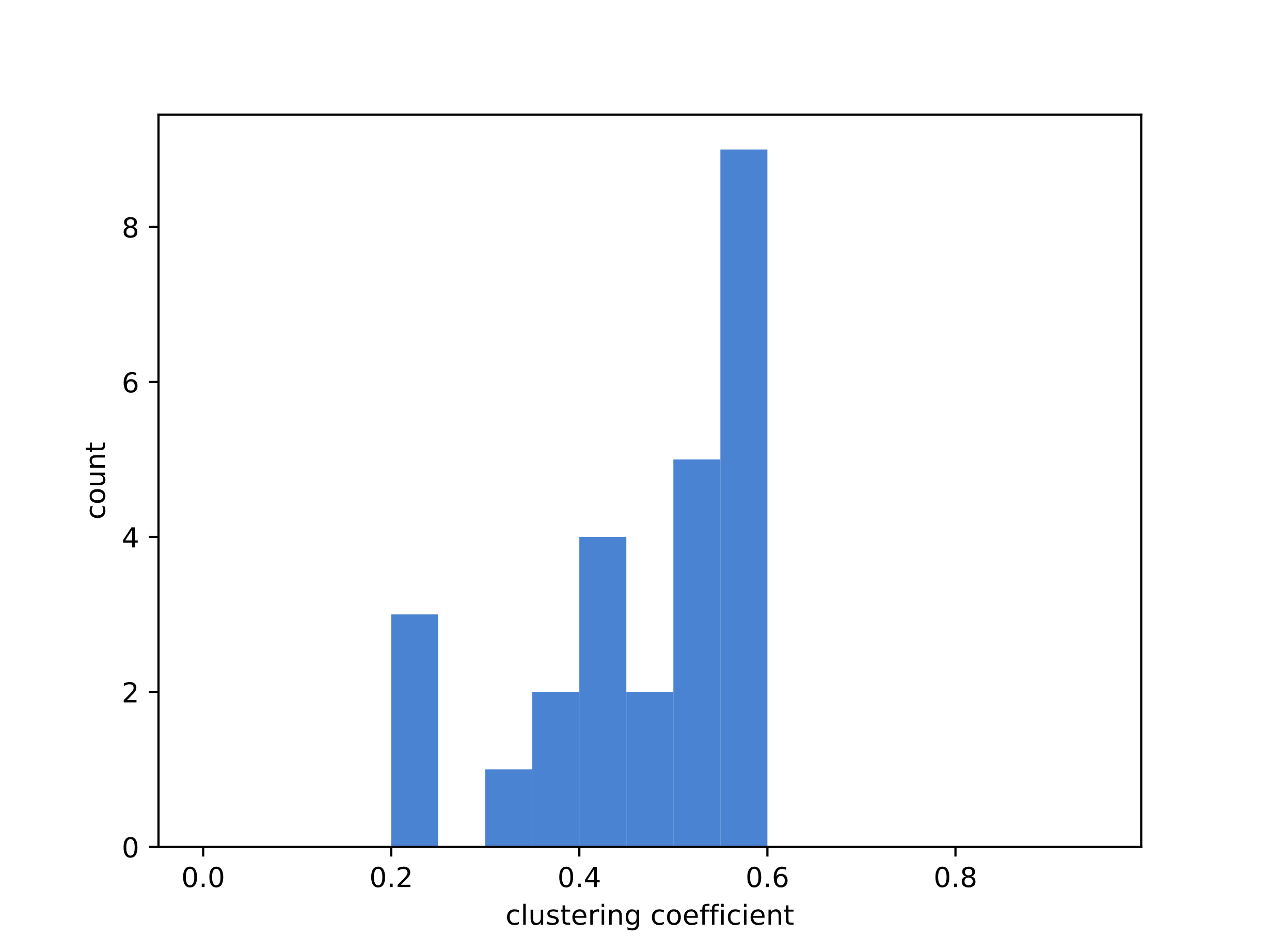

Figure 2 Distribution of the clustering coefficient for all CIR type modules.

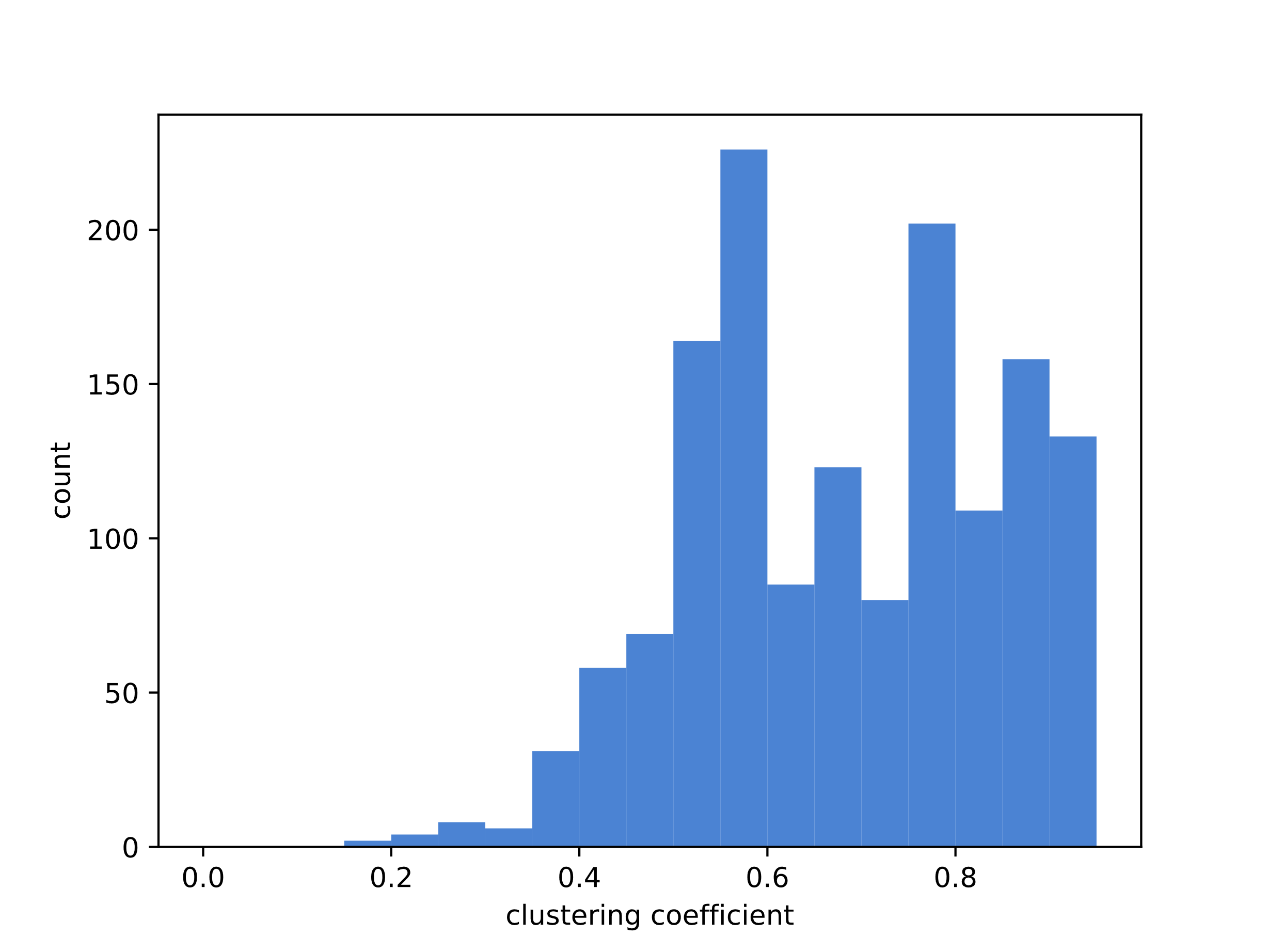

Figure 3 Distribution of the clustering coefficient for all COM type modules.

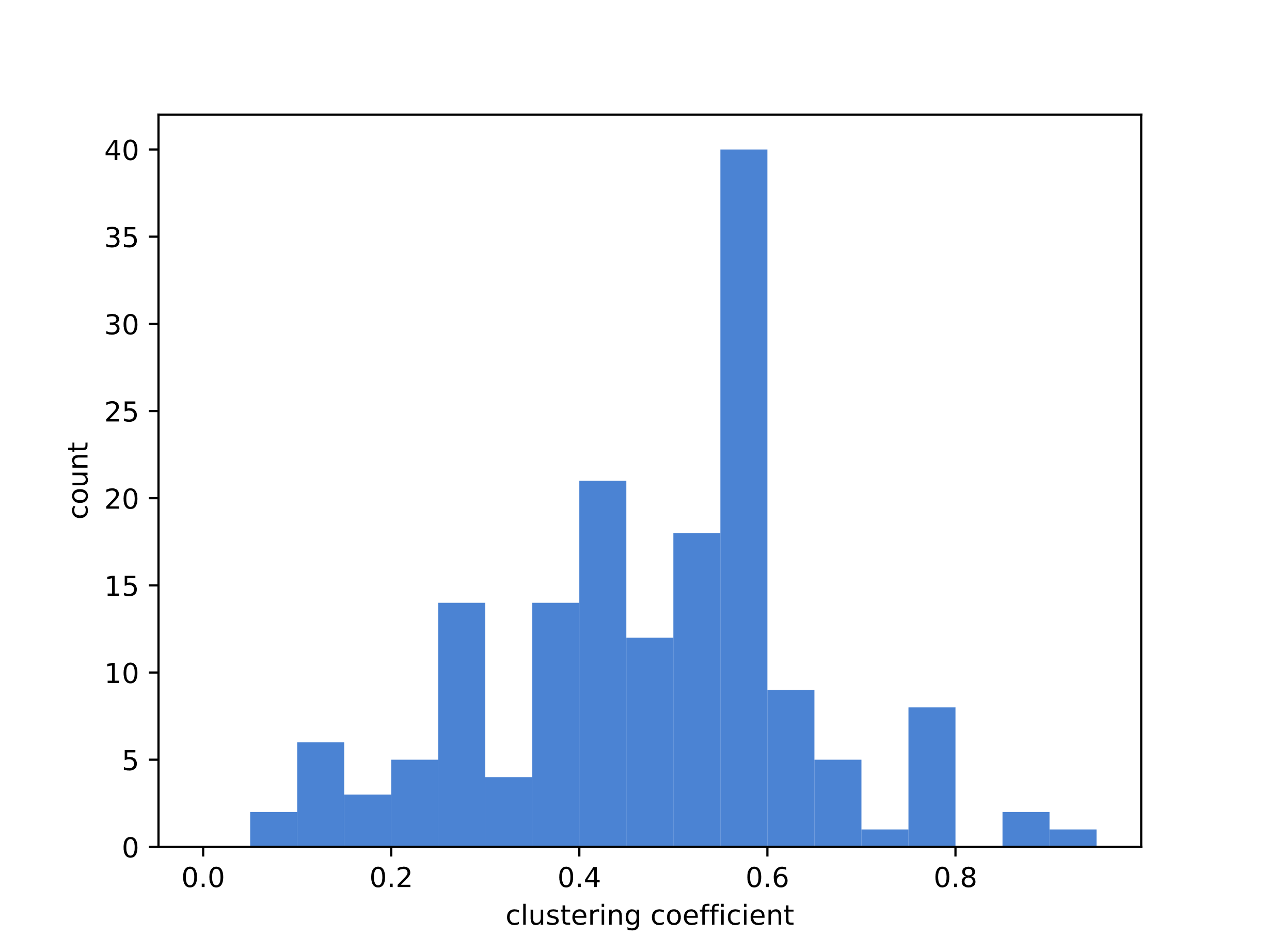

Figure 4 Distribution of the clustering coefficient for all COP type modules.

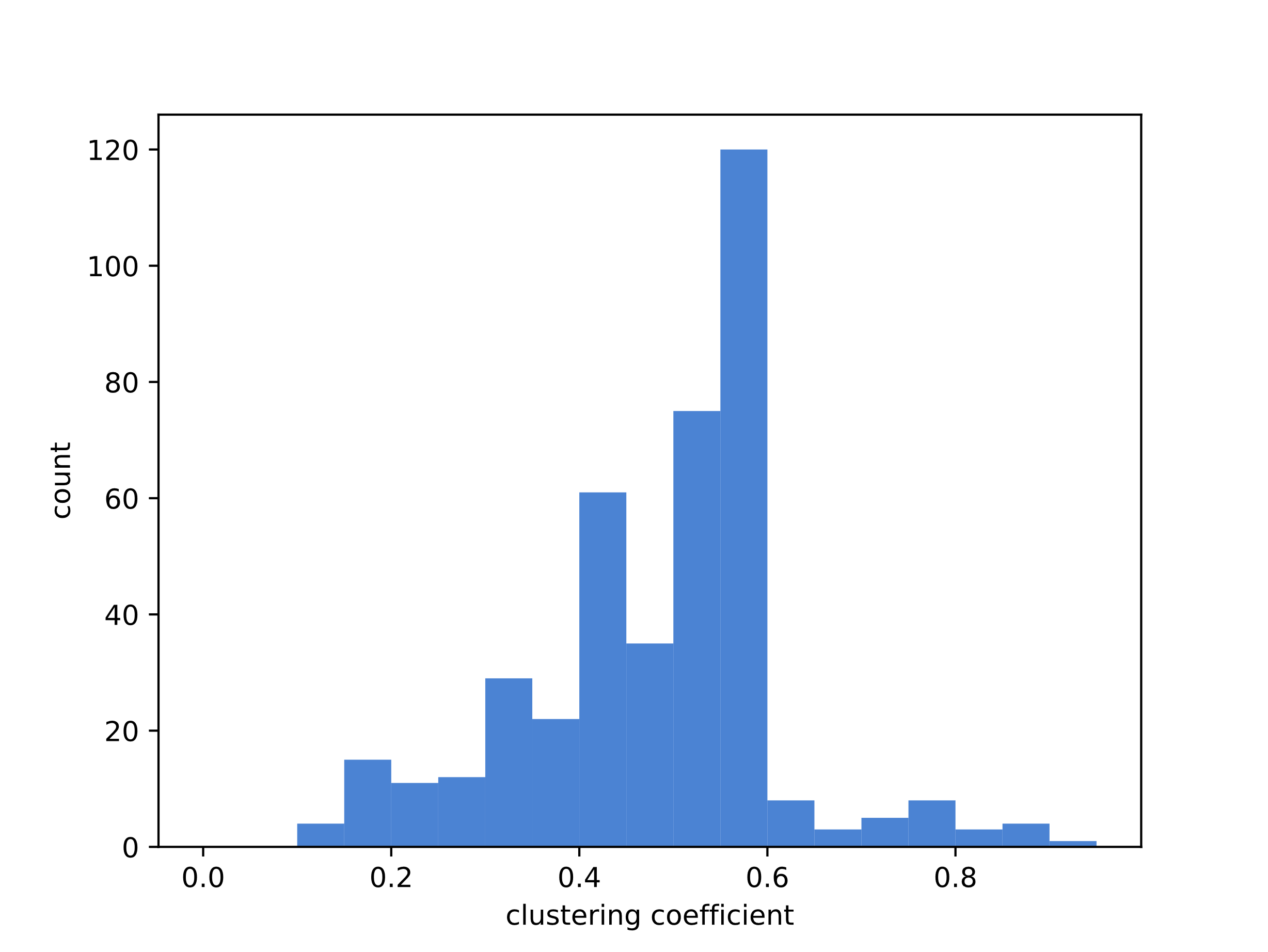

Figure 5 Distribution of the clustering coefficient for all COR type modules.

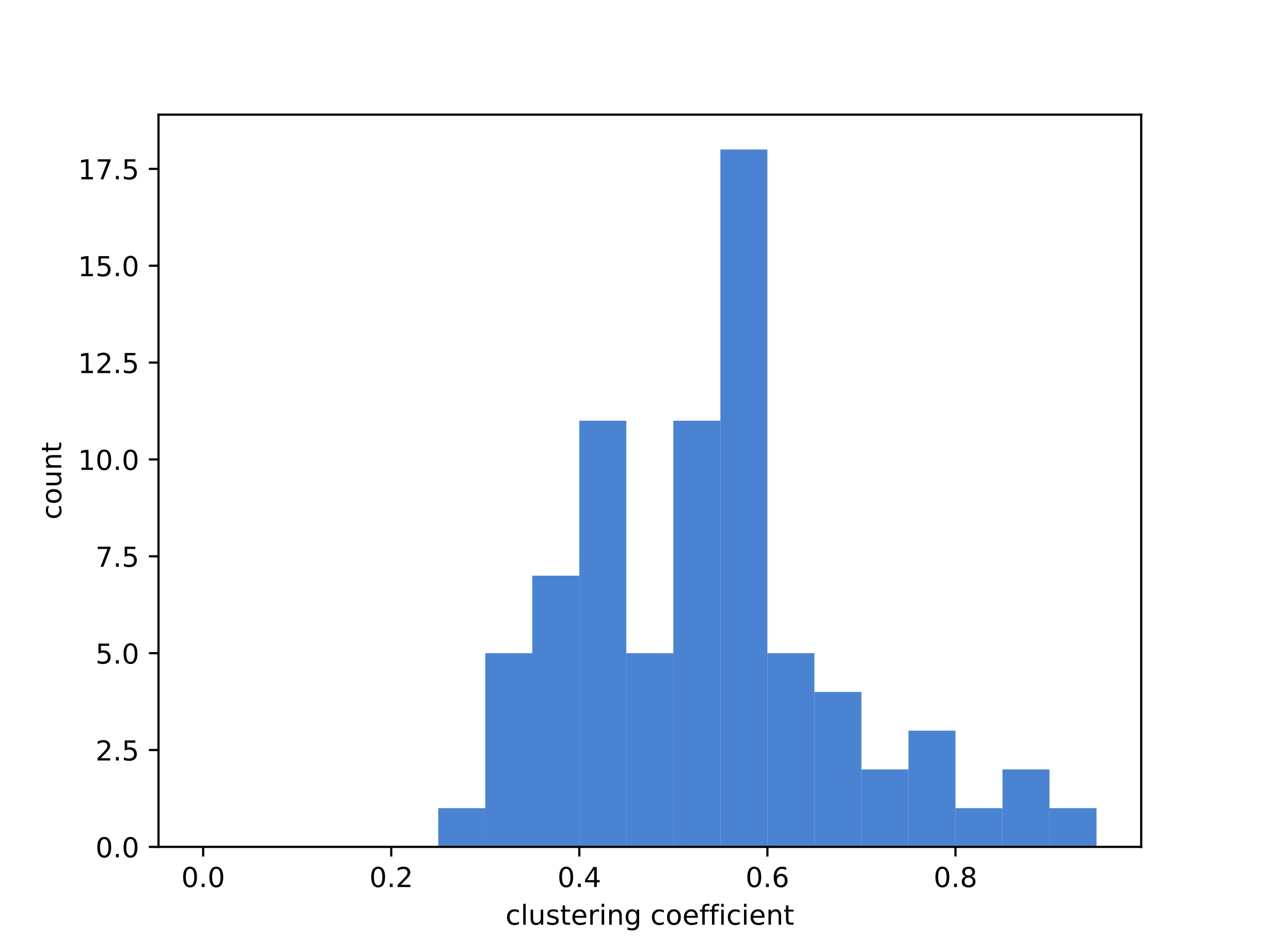

Figure 6 Distribution of the clustering coefficient for all FB2U type modules.

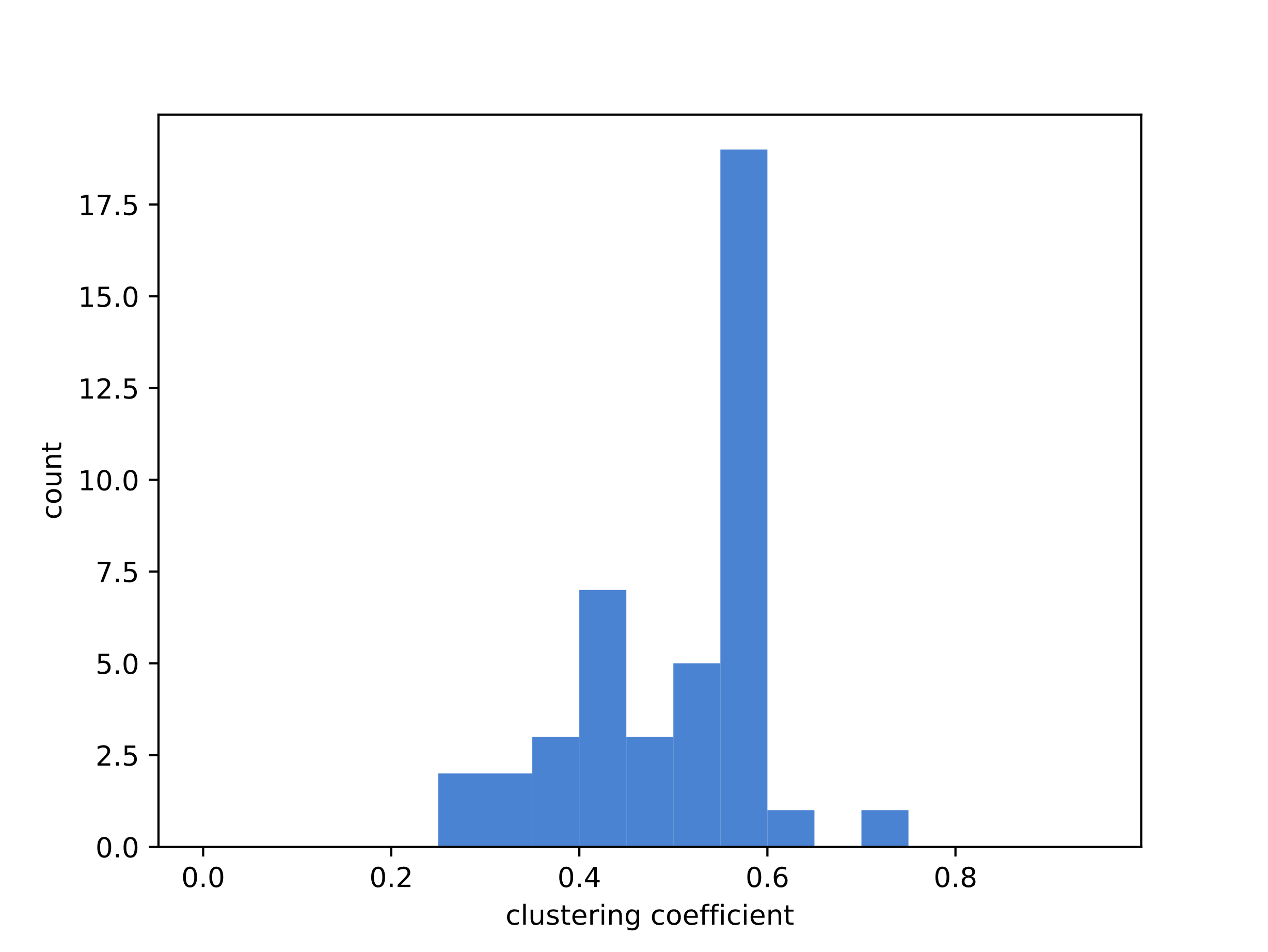

Figure 7 Distribution of the clustering coefficient for all FBU type modules.

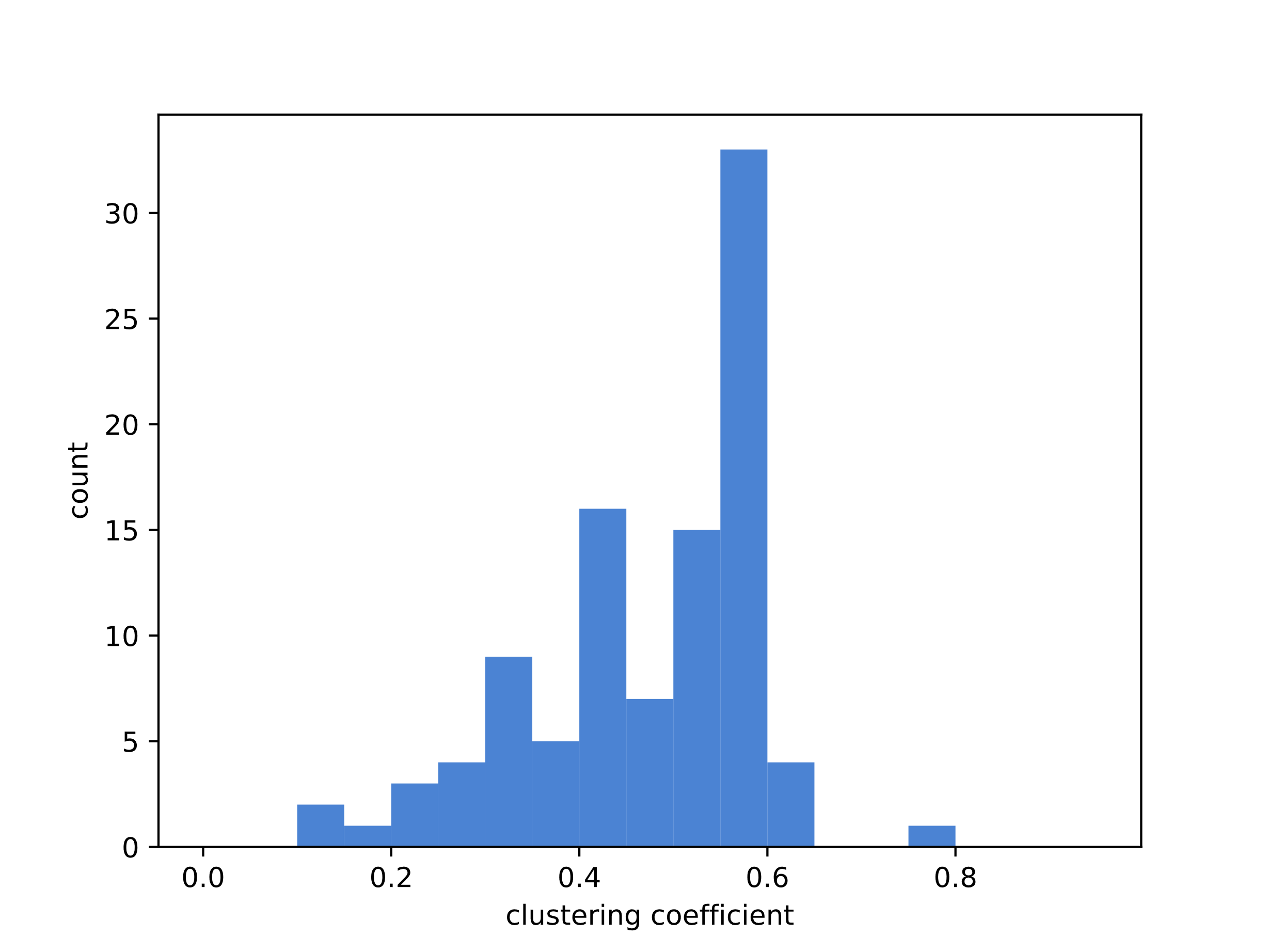

Figure 8 Distribution of the clustering coefficient for all FFL type modules.

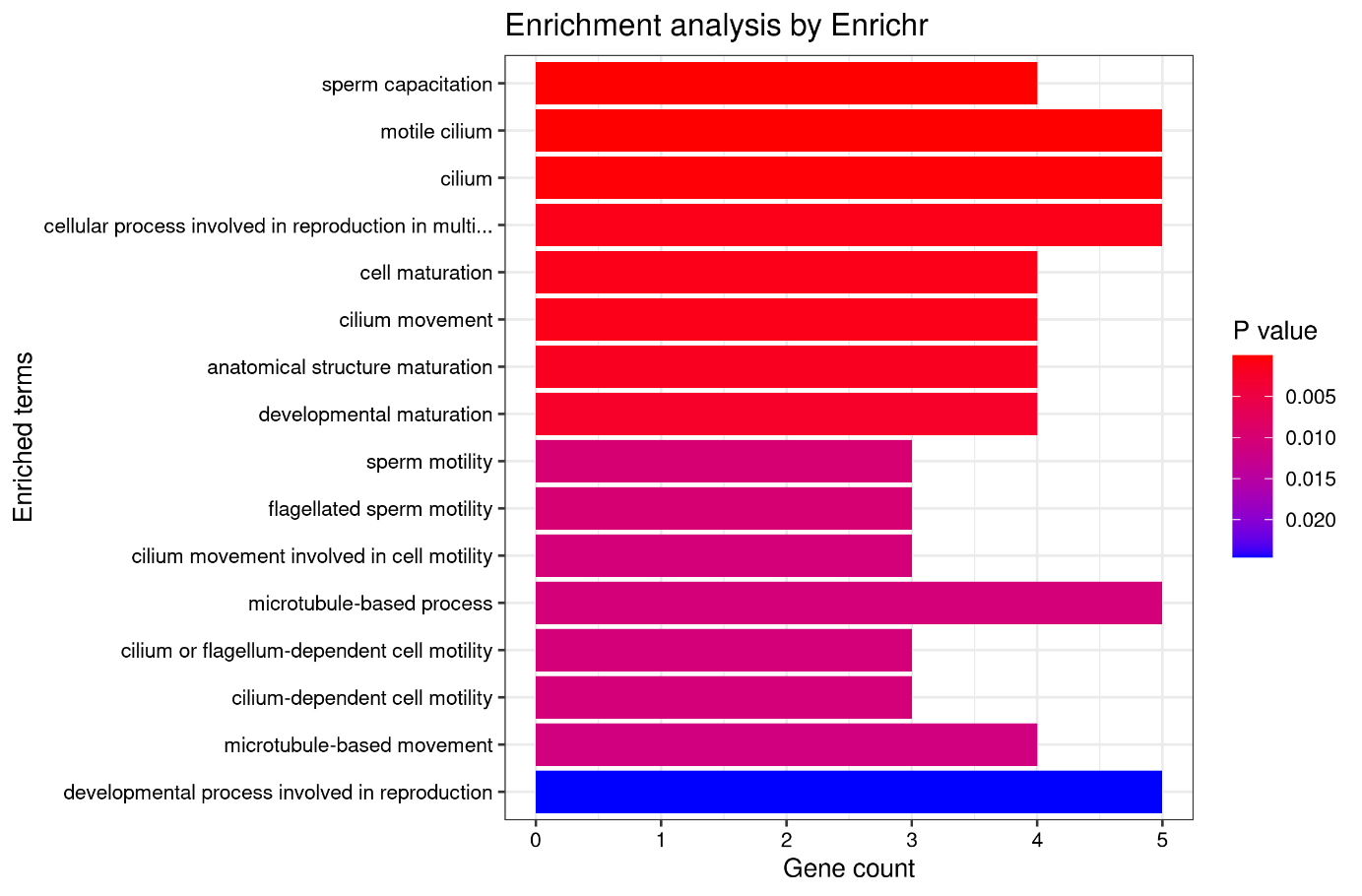

Figure 9 Enrichment of genes in the ALL_331 module. Genes were ranked by p-value.
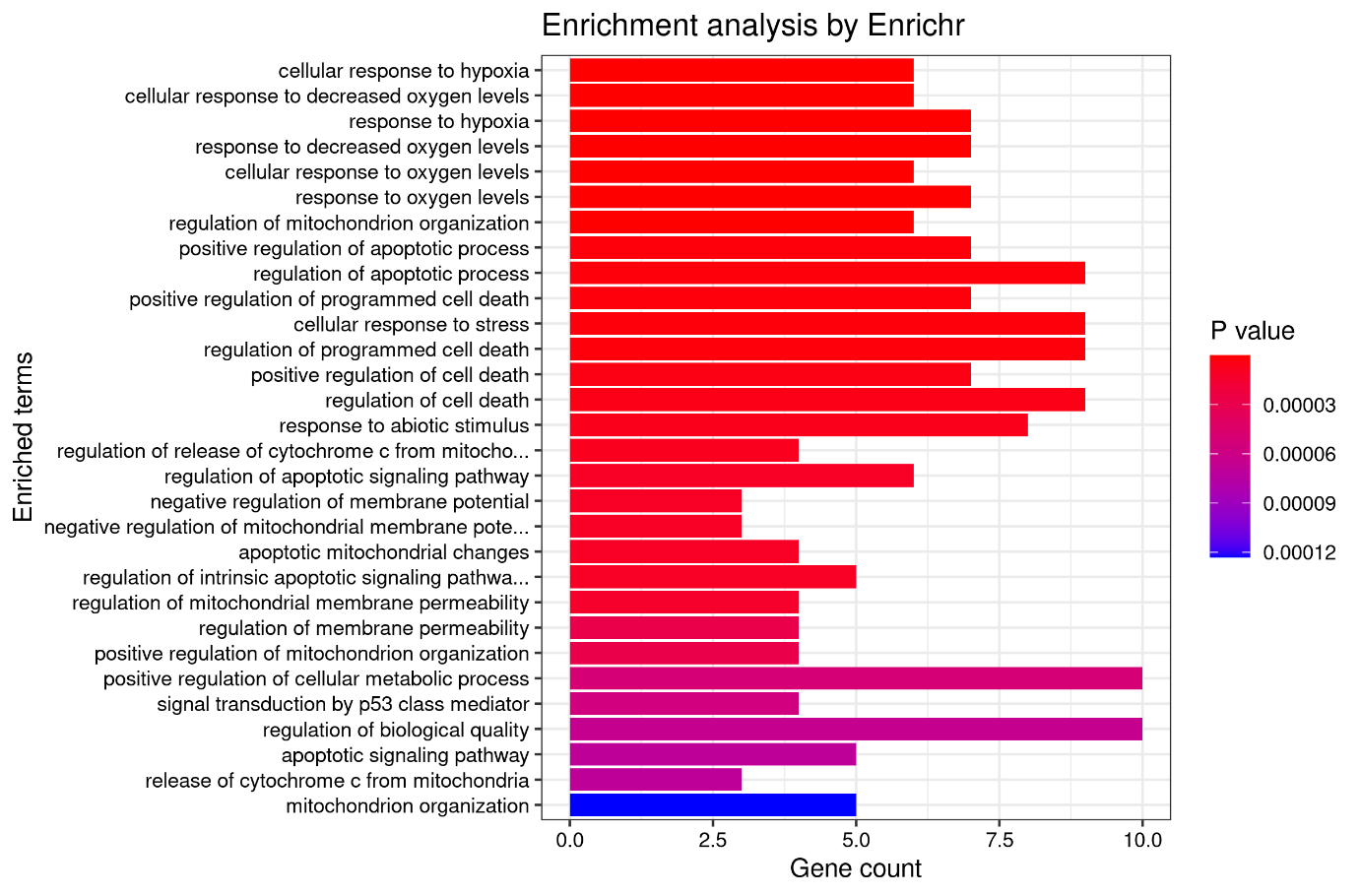

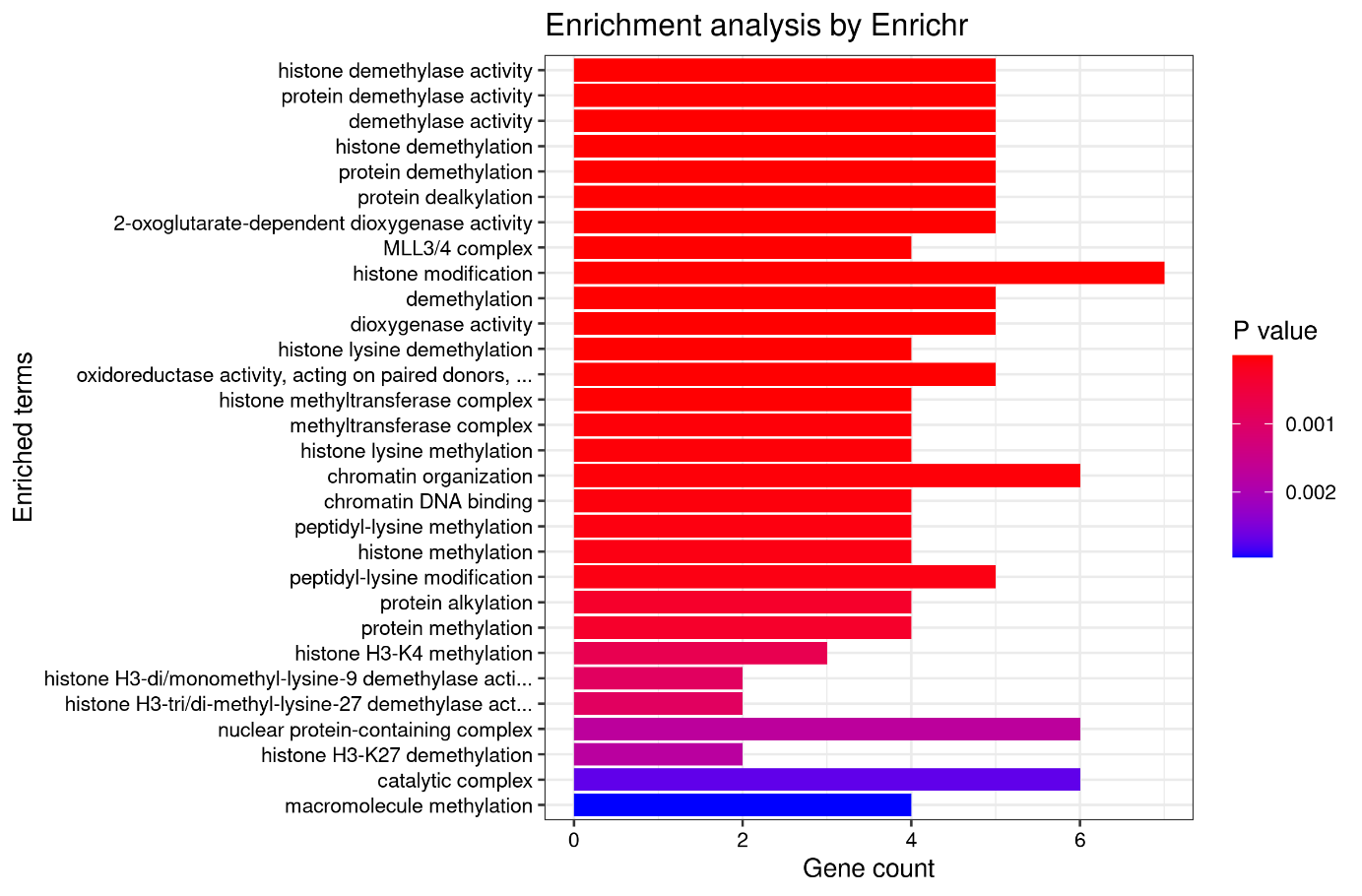

Figure 10 Enrichment of genes in the ALL_1135 Module. Genes were ranked by p-value.

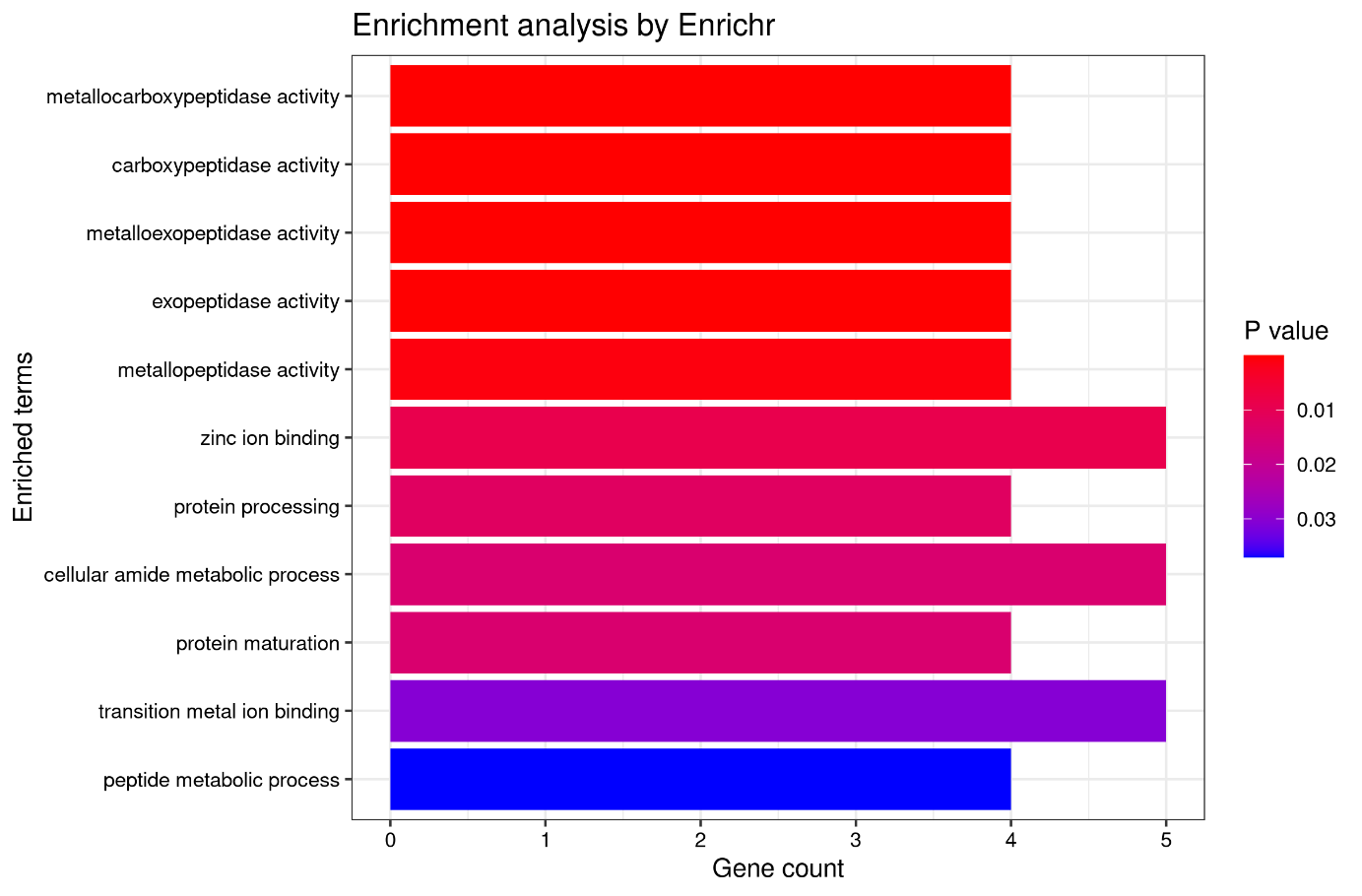

Figure 11 Enrichment of genes in the ALL_1153 module. Genes were ranked by p-value.

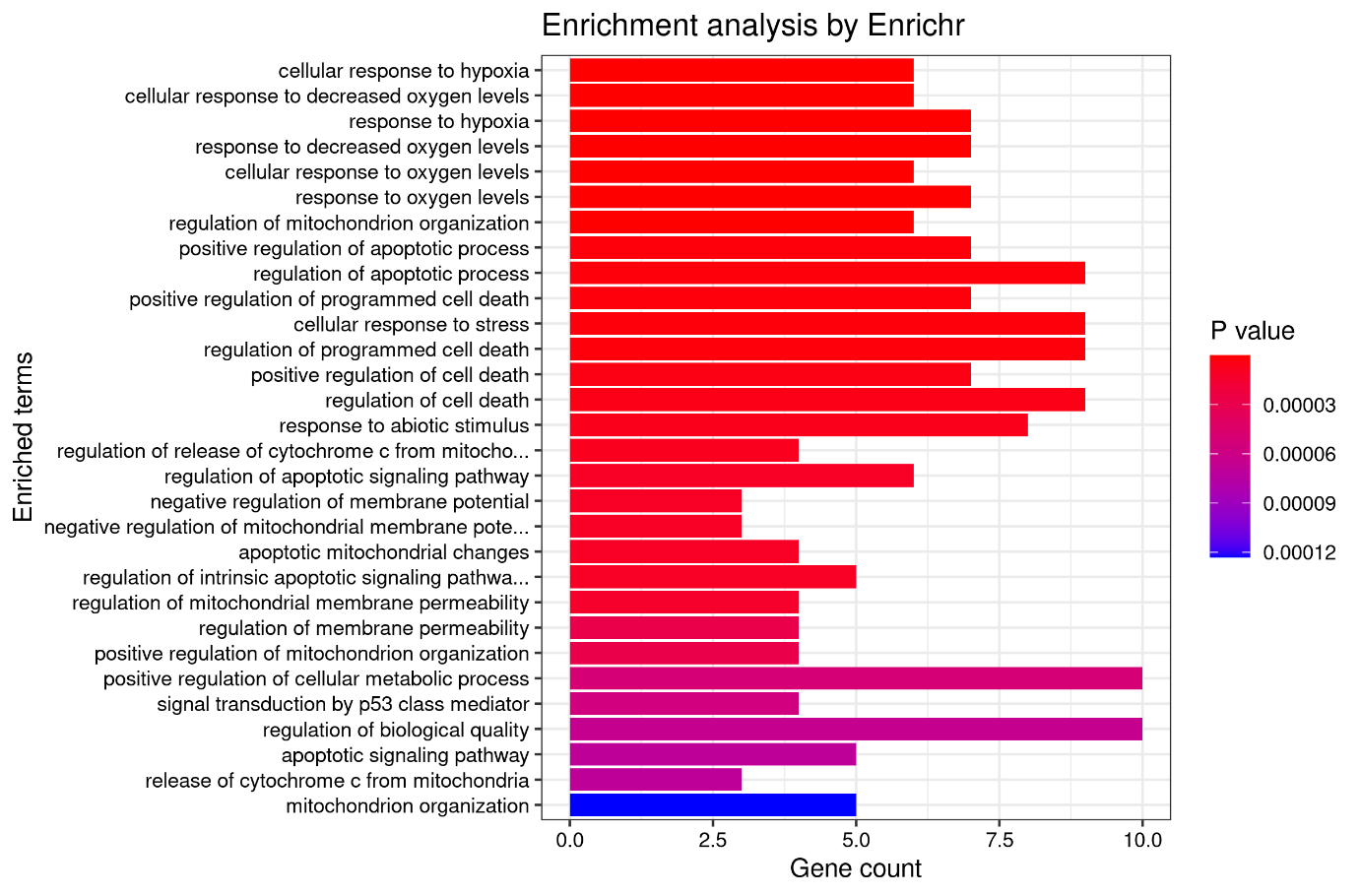

Figure 12 Enrichment of genes in the ALL_1753 module. Genes were ranked by p-value.

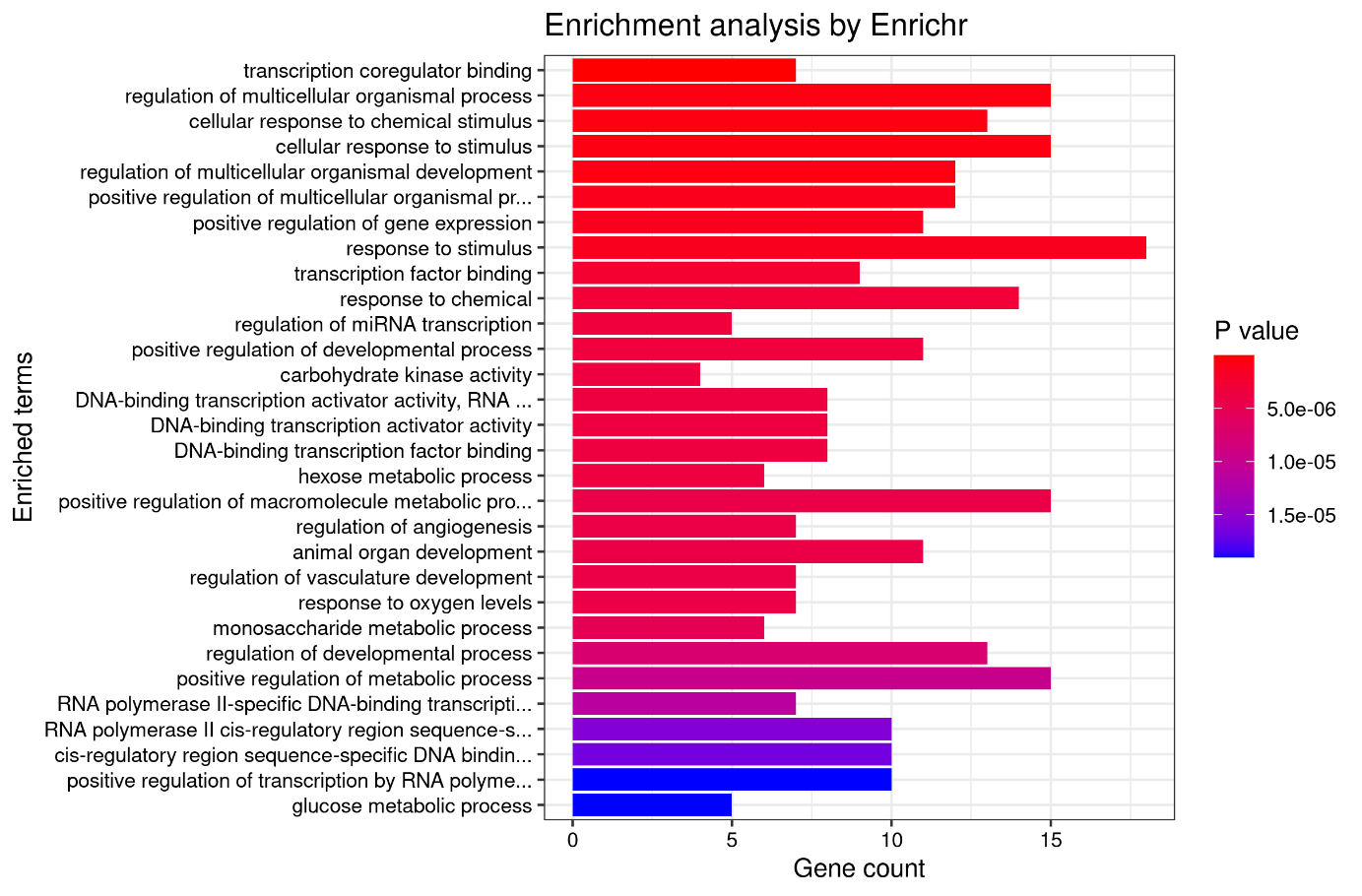

Figure 13 Enrichment of genes in the ALL_2093 module. Genes were ranked by p-value.

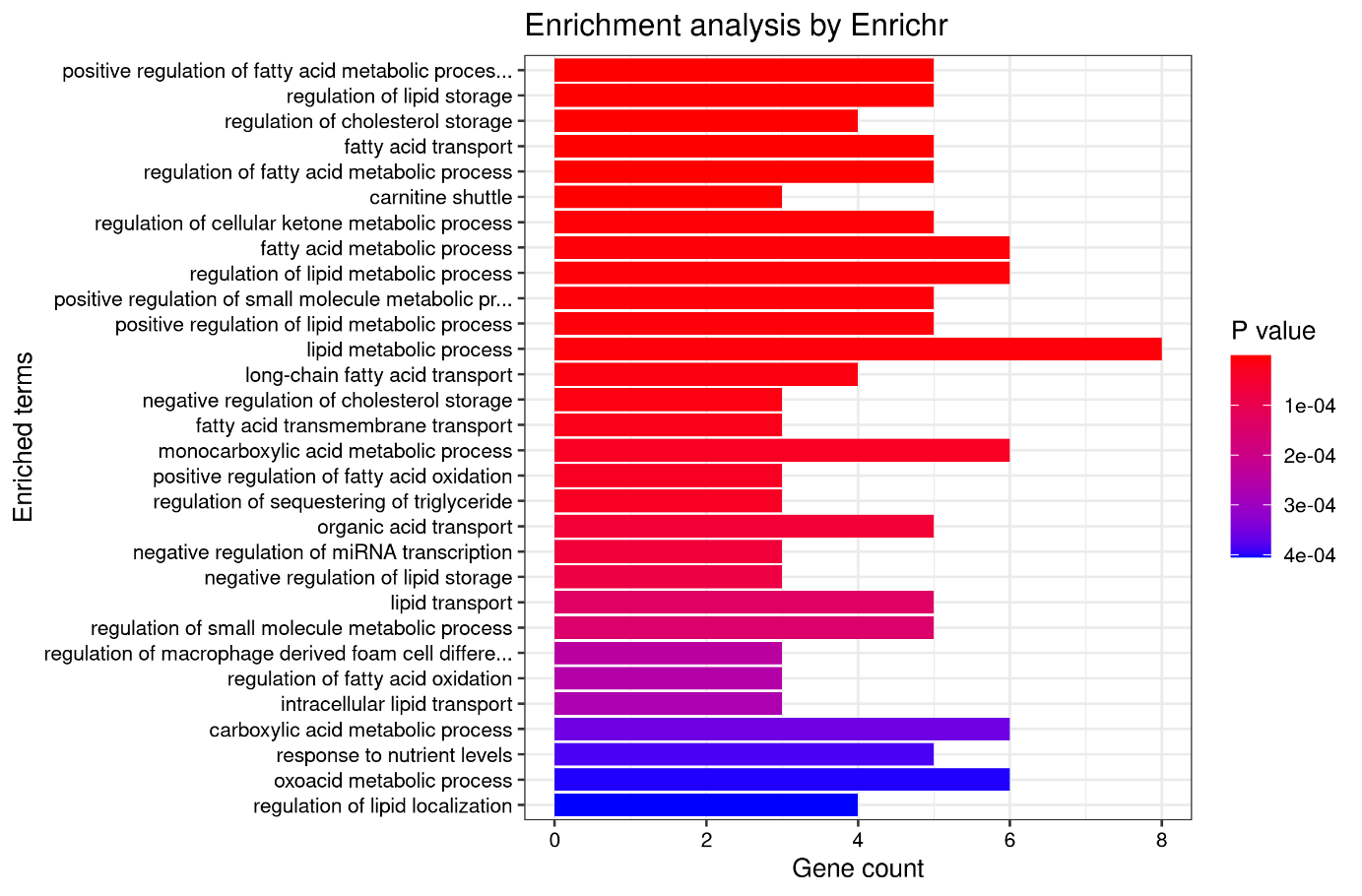

Figure 14 Enrichment of genes in the ALL_2269 module. Genes were ranked by p-value.

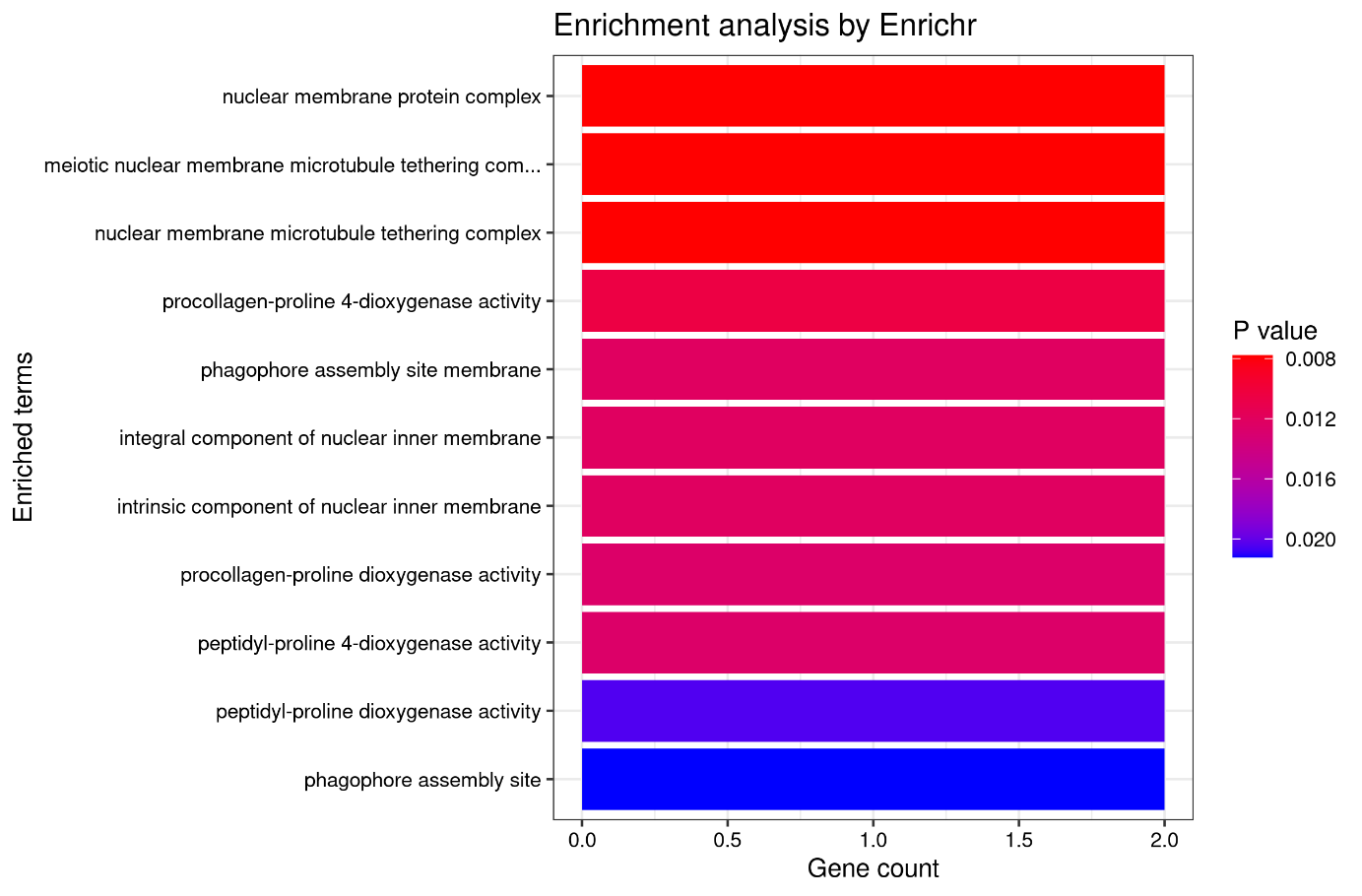

Figure 15 Enrichment of genes in the ALL_2896 module. Genes were ranked by p-value.

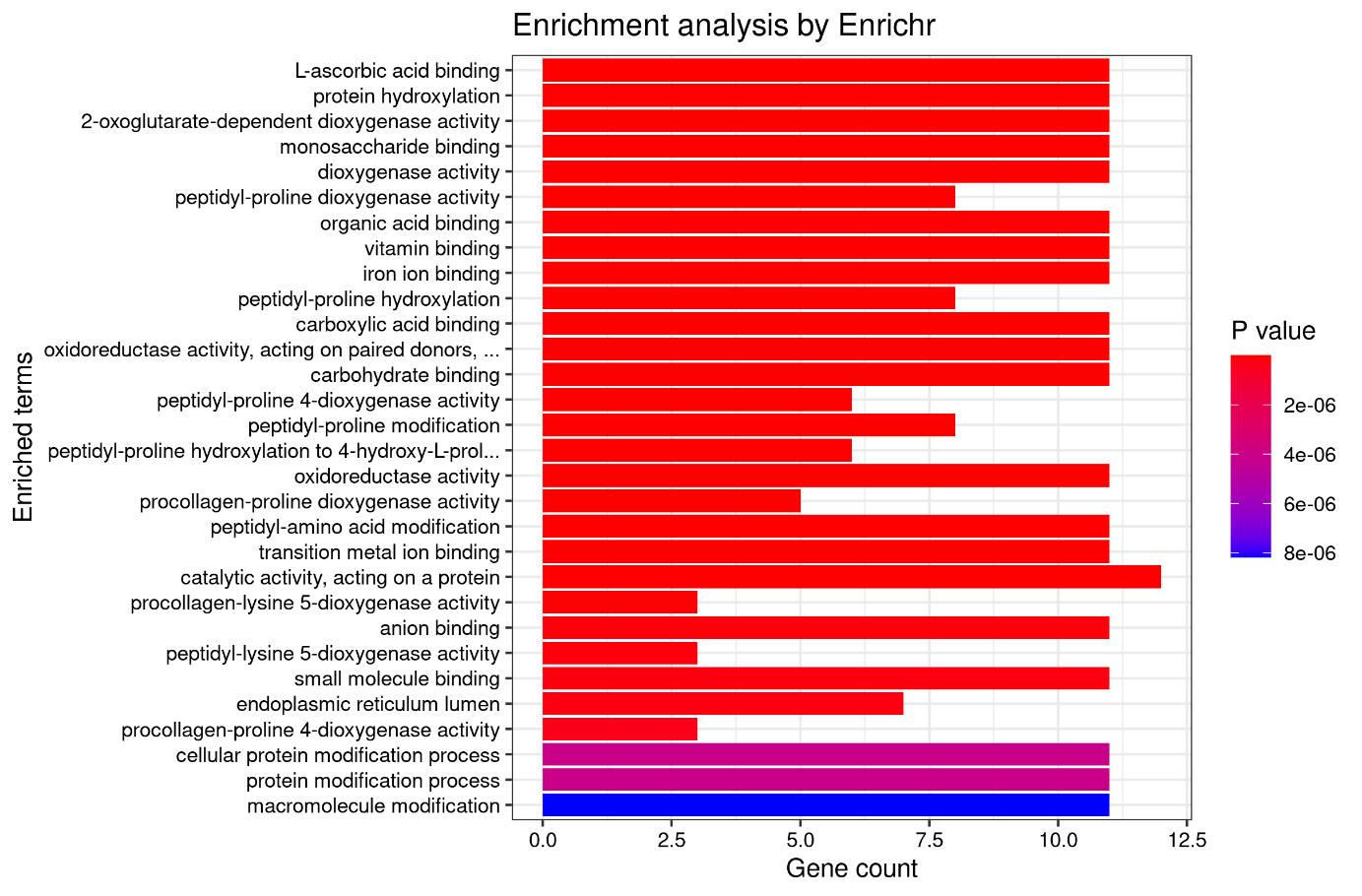

Figure 16 Enrichment of genes in the COM_256 module. Genes were ranked by p-value.

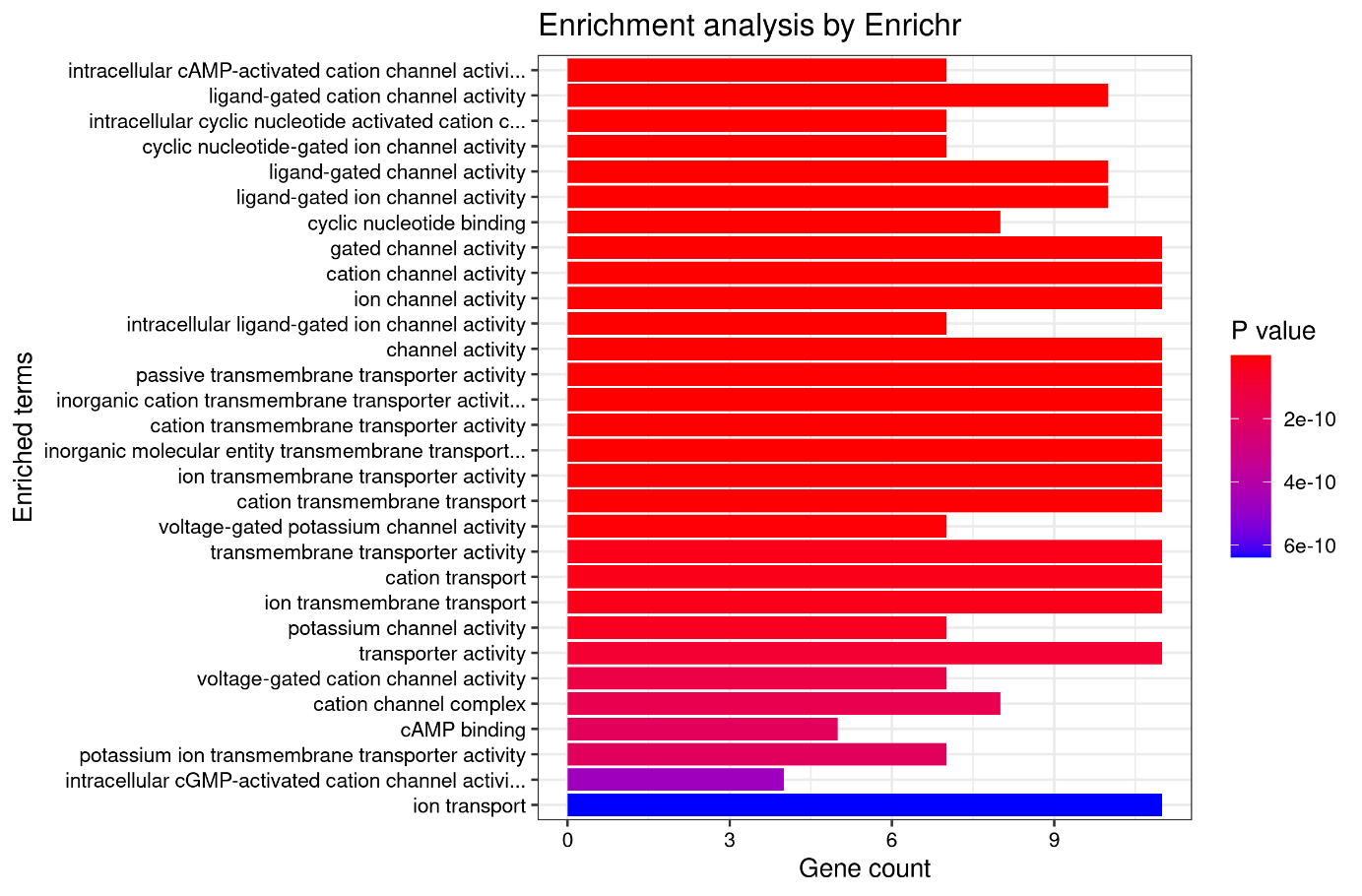

Figure 17 Enrichment of genes in the COM_667 module. Genes were ranked by p-value.

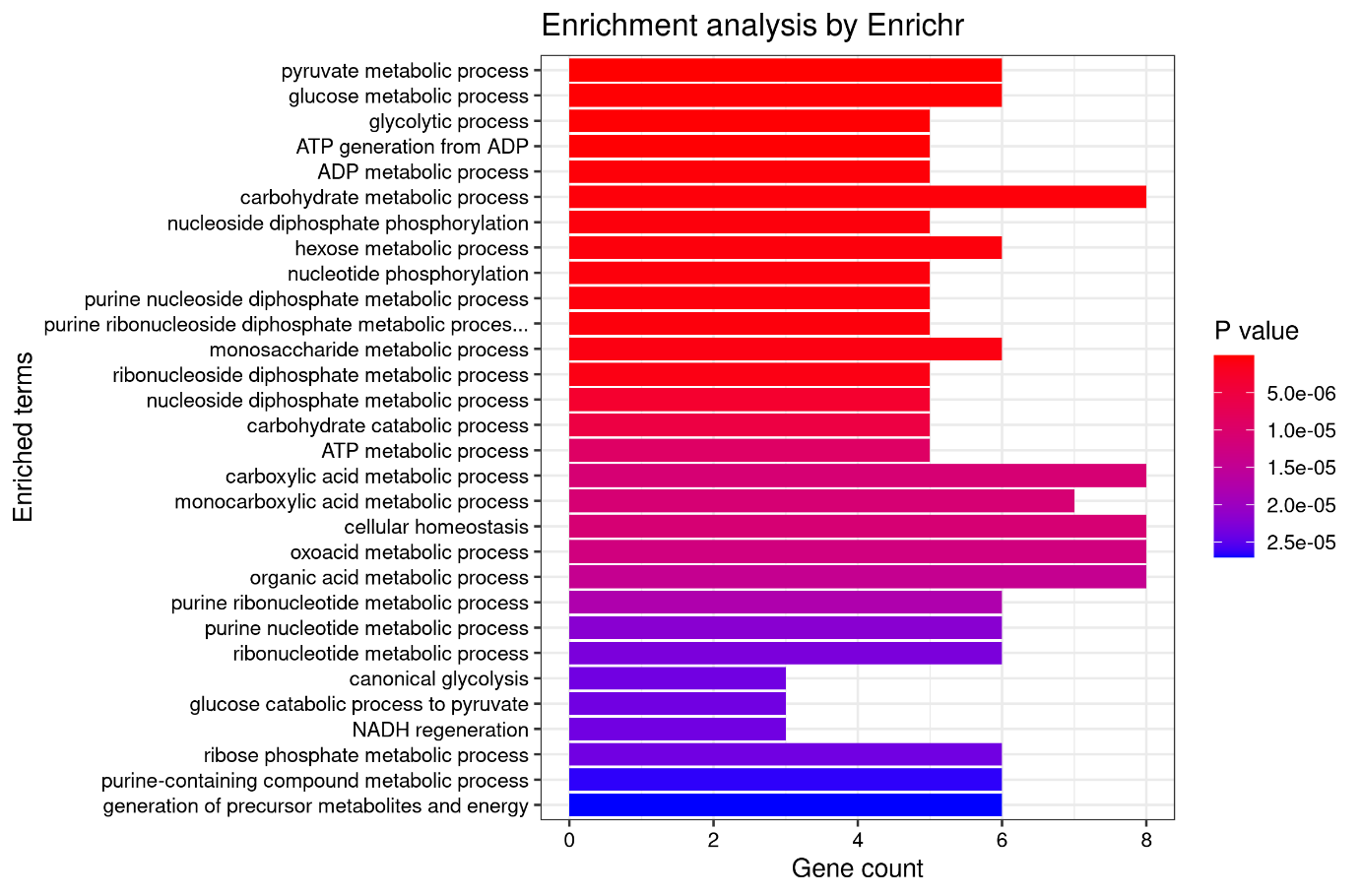

Figure 18 Enrichment of genes in the COR_347 module. Genes were ranked by p-value.

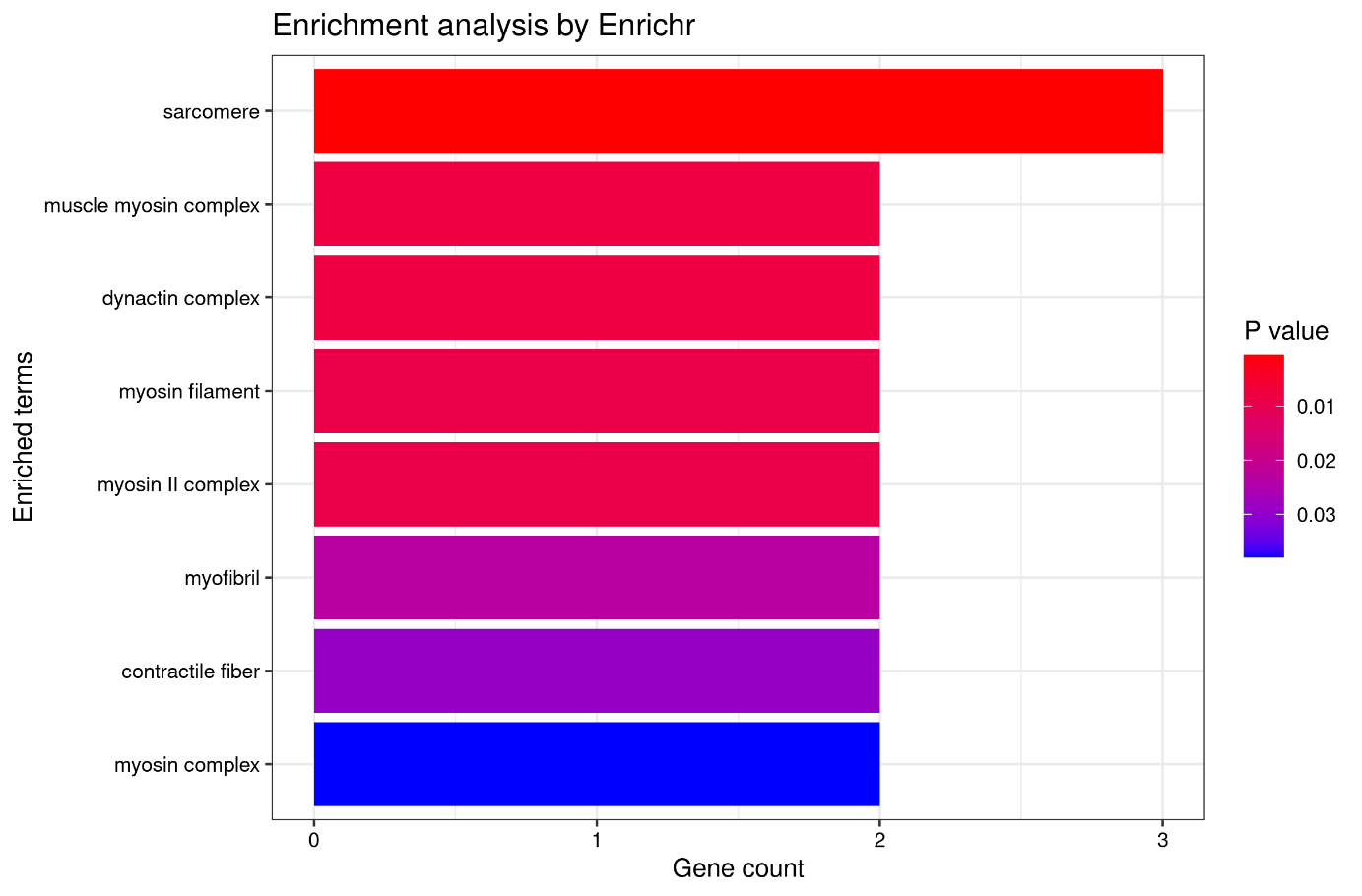

Figure 19 Enrichment of genes in the FB2U_87 module. Genes were ranked by p-value.

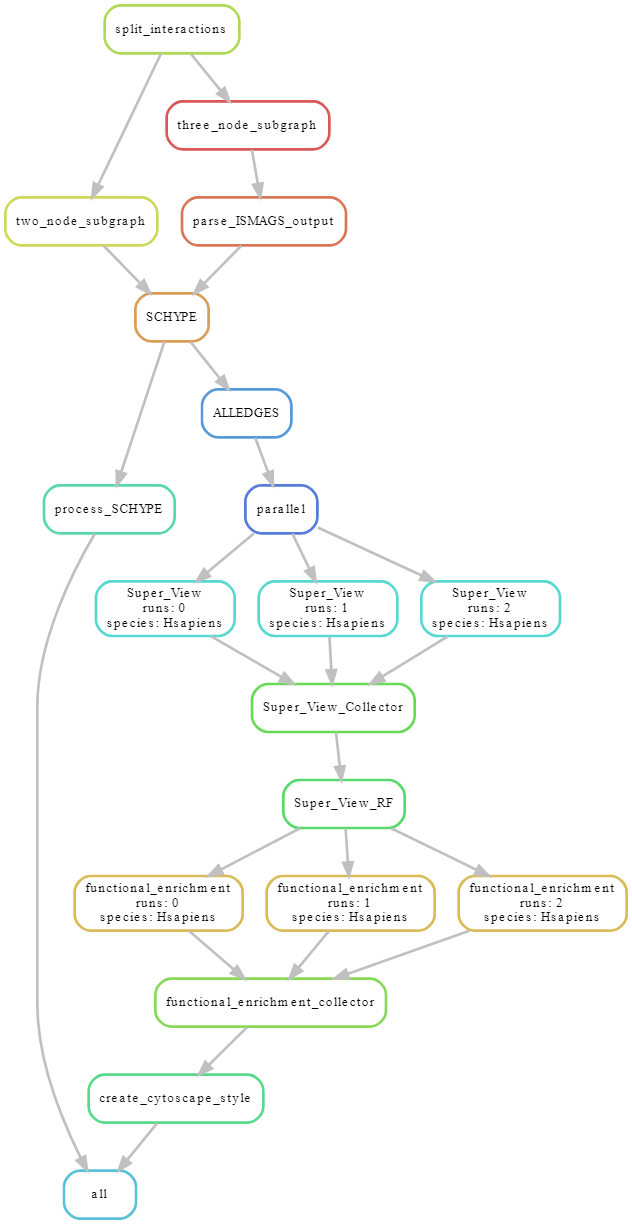

Figure 20 Schematic overview of the Snakemake pipeline in an example run on three different cores

1. Xie Z, Bailey A, Kuleshov MV, Clarke DJB, Evangelista JE, Jenkins SL, et al. Gene Set Knowledge Discovery with Enrichr. Current Protocols. 2021;1:e90.
